# Spatial genomics of the cardiac sarcomere

**DOI:** 10.1101/2025.10.07.680947

**Authors:** Christian Michael Cadisch, Victoria Parikh, Megan Grove, Aditya Singh, Dominic Abrams, Neal Lakdawala, Carolyn Ho, Erin Miller, Thomas Ryan, Joseph Rossano, Kimberly Y. Lin, Anna Axelsson Raja, Henning Bundgaard, Michelle Michels, Peter Paul Zwetsloot, Jodie Ingles, Alexandre Pereira, Adam Helms, Sara Saberi, Lia Crotti, Anjali Owens, Sharlene M. Day, Belinda Gray, John C. Stendahl, Rachel Lampert, Iacopo Olivotto, Alessia Argiro, Niccolo Maurizi, George Powell, James Ware, Masataka Kawana, Euan A. Ashley

**Author notes:** **Correspondence**, Euan A. Ashley, 300 Pasteur Drive, Stanford, California 94305, United States.

## Abstract

**Background:** Structure–function mapping of proteins has improved our understanding of disease mechanisms and protein domains, while human population genetics has provided a window into variational tolerance that can be modeled at a structural level. To reveal new insight into the cardiac sarcomere, the motor unit of the heart, we developed an analytical framework that integrates large-scale genetic and structural data.

**Methods:** Data from more than 17,000 patients with hypertrophic cardiomyopathy (HCM) were first analyzed and contrasted with two population-scale genomic databases comprising over 800,000 individuals. Building on this clinical and population-level analysis, we developed an integrative analytical framework that supplements observed variants with selected AlphaMissense-predicted missense substitutions and applies three-dimensional integrative spatial scanning across multiple structural models of cardiac motor proteins.

**Results:** This integrative approach identified both recognized and novel regions of structural variant intolerance across critical genes of the cardiac sarcomere and provided new insight into protein function. Structural relevance of variant enrichment was interpreted in the context of sarcomere organization and destabilization of sequestered myosin leading to hypercontractility in HCM, incorporating the recently defined high-resolution structure of the human cardiac myosin filament. These findings were further assessed through analyses of pathogenic variant class enrichment and demonstrated novel associations with earlier onset of disease in a large clinical cohort.

**Conclusions:** Our study provides a multidimensional framework that integrates structural, genomic, and modeling data to advance understanding of the cardiac sarcomere and the molecular mechanisms underlying hypertrophic cardiomyopathy, highlighting previously unrecognized regions for functional investigation and clinical evaluation.

## Introduction

Structure-function mapping of proteins has improved our understanding of disease mechanisms and critical protein domains. Specifically, mapping the population-level disease association of variants can help to identify regions intolerant to variation that can be inferred to have disproportionate functional importance. We have previously shown that extension of these comparisons to three-dimensional space can greatly increase the information gleaned about proteins of the cardiac sarcomere, and variants that cause hypertrophic cardiomyopathy (HCM) [1], [2]. HCM is a genetic disease of the heart muscle characterized by left ventricular hypercontractility and hypertrophy, and affects ∼1 in 500 individuals [3]. The clinical course of HCM is notably variable, with some patients experiencing minimal symptoms while others develop severe complications including arrhythmia, heart failure, or sudden cardiac death [3].

HCM is primarily a disease of the cardiac sarcomere. Variants in over a dozen genes encoding sarcomere-associated proteins are known to cause HCM, among these, variants in the *MYH7* gene, encoding the β-cardiac myosin heavy chain, and *MYBPC3*, encoding myosin-binding protein C, are the most prevalent causes [4]. The beta myosin heavy chain protein includes a globular head domain where ATP binding, hydrolysis, and actin interaction occur, followed by a lever arm and a long alpha-helical coiled-coil rod domain. Over 200 disease-associated variants, primarily missense, have been identified in *MYH7*, with a significant concentration in the motor domain of the head region, which ClinGen (a National Institute of Health funded central resource for variant clinical relevance) recognizes as a pathogenic hotspot [5]. Past research has identified the converter domain and the myosin mesa, a relatively flat surface on the myosin motor domain composed of several myosin subdomains that is conserved in all cardiac myosin from all species, as enriched for variants associated with HCM with a more severe prognosis and earlier disease onset [1], [6], [7]. The clinical genetics and functional analysis of these variants led to significant improvements in understanding the structure-function relationships underlying the hypercontractility observed in HCM, and informed design of several putative therapies now commercially approved or pending Food and Drug Administration (FDA) review [8], [9], [10]. Early experience with these precision therapies shows a significant reduction in symptoms and biomarkers of disease severity [9], [10].

In contrast to *MYH7*, mapping of disease-observed variants to known or predicted 3D protein structures of other sarcomeric proteins has not been performed. While other common sarcomeric genes known to cause HCM, such as *MYBPC3*, have had several enriched regions identified by 2D mapping [11], most *MYBPC3* disease-observed variants are truncating rather than missense variants. Similarly, disease-observed variants in genes encoding the troponin complex proteins (*TNNT2*, *TNNI3*); and the myosin light chains (*MYL2*, *MYL3*) are generally less common [12]. The lack of structure-function modeling integrating clinical data for these genes may stem from the lower prevalence of disease-observed variants and the resulting limitations in statistical power. Previous attempts to identify disease-associated regions have employed various approaches, from spatial scan statistics [1] to generalized additive modeling [11], but have been limited by the available data and computational tools. As *MYH7* is a well-studied gene with prior 3D analyses and functional validation, we first re-examined it to validate our methodology and determine whether additional data would reveal previously unrecognized structural insights [8].

Recent technological advances have dramatically enhanced our ability to study structure-function relationships at the level of single amino acid substitutions in the sarcomere. Numerous large-scale biobanks with genetic and imaging data from both disease-affected and general populations have enabled the compilation of extensive datasets of human genetic variation. Furthermore, the introduction of transformer-based computational models like AlphaFold (and the related AlphaMissense predictor) has significantly improved our ability to predict variant pathogenicity [13] and protein structure [14]. In addition, advancement in structural biology and particularly cryogenic electron microscopy (cryo-EM) has enabled us to obtain high-resolution images of human sarcomere structure and individual proteins such as myosin, which would further improve the understanding of the inter- and intra-molecular interactions that are critical in defining sarcomere function [15], [16], [17], [18], [19]. Combining clinical genetics and structural biology holds high promise for fulfilling the current unmet need for improving variant assessment in daily practice where clinicians encounter a large number of variants of unknown significance.We hypothesized that integrating these advanced computational tools with clinical and population genetic databases could reveal novel insights into the structure-function relationships within key sarcomeric genes and identify previously unknown variant enrichments. In this study, we combine exomic data representing 814,392 individuals with the Sarcomeric Human Cardiomyopathy Registry (SHaRe), a HCM clinical database of over 17,000 HCM patients, and AlphaMissense pathogenicity predictions. Using a spatial scan statistic approach, we investigated regions of disease enrichment in the 3D protein structures of MYH7, MYBPC3, TNNT2, TNNI3, MYL2 and MYL3. Finally, we assessed whether identified regions were enriched with variants classified as pathogenic or likely pathogenic in ClinVar, and explored phenotypic differences across the cohorts, focusing on the age of disease onset.

## Materials and Methods

### Analytical overview

We first compared unique variants observed in patients with HCM in SHaRe with variants observed in population datasets. We then repeated the analysis after supplementing these observations with selected high- and low-scoring AlphaMissense substitutions. Variants were mapped to three-dimensional protein models, and a spatial scan identified regions with a higher proportion of HCM-cohort variants, or of the corresponding augmented class, than the rest of the protein. We assessed the identified regions using ClinVar classifications, clinical age-of-onset data and an external HCM dataset. Detailed structural modeling procedures and model comparisons are provided in the Supplemental Methods.

### Datasets

We have used an extensive dataset of genetic information, combining clinical data, exonic genetic data and pathogenicity prediction scores (see Table 1).

**Table 1.** Overview of genetic variation and in silico pathogenicity predictions across six key sarcomeric genes. This table highlights the scale of the study, combining real-world clinical and population data from 800,000 individuals (gnomAD and UKBB) and 12,326 phenotyped HCM patients (SHaRe registry), alongside over 75,000 pathogenicity predictions scores to support structure-function analysis.

| **Gene** | **No. Variants in Population Datasets** | **No. Patients with Variants in SHaRe** | **No. Variants observed in SHaRe** | **No. Pathogenicity Predictions** |
|---|---|---|---|---|
| *MYH7* | 1,893 | 1,332 | 363 | 38,700 |
| *MYBPC3* | 1,852 | 2,523 | 522 | 20,480 |
| *MYL2* | 274 | 103 | 30 | 3,320 |
| *MYL3* | 231 | 58 | 22 | 3,900 |
| *TNNT2* | 392 | 218 | 63 | 5,960 |
| *TNNI3* | 336 | 184 | 56 | 4,200 |

#### Sarcomeric Human Cardiomyopathy Registry (SHaRe)

The SHaRe is a collaborative initiative that aggregates anonymized patient data from multiple research institutions to advance the study of cardiomyopathies. It includes information from patients diagnosed with HCM, DCM, and arrhythmogenic cardiomyopathy. The registry contains genetic, clinical, and demographic information from over 17,000 patients, of which 12,326 were diagnosed with HCM. Of these, 4,427 patients have genetic variants within the six genes analyzed in this research, totaling 1,056 unique variants.

#### UK Biobank (UKBB)

The UK Biobank is a comprehensive repository of biomedical data and research materials. It contains anonymized information on genetics, lifestyle habits, and health outcomes, along with biological samples, collected from 500,000 individuals across the United Kingdom. Unlike disease-focused databases such as SHaRe, UKBB is curated from the general population and not enriched for people with specific conditions. The effort results in 346,977 patients with variants in the genes of interest for our study, yielding a total of 1,479 unique variants without applying and filtering based on minor allele frequency MAF.

#### Genome Aggregation Database (gnomAD)

The Genome Aggregation Database, originally established as the Exome Aggregation Consortium (ExAC), represents a global collaboration among researchers committed to sharing data from exome and genome sequencing projects. Similar to the UKBB, gnomAD compiles data from the general population. In its latest release (v4), gnomAD has expanded its database to include 730,947 exomes of which 416,555 originate from the UK Biobank. Among these, 586,403 patients have variants in the sarcomeric genes we focused on, amounting to 4,620 unique observed variants without applying any filtering based on MAF.

#### AlphaMissense

AlphaMissense is a transformer-based model developed to predict the pathogenicity of missense variants in the human genome [13]. For each gene, the model creates pathogenicity predictions for the 19 possible missense variants at each amino acid position. Within the six genes we studied, this results in a total of 77,482 pathogenicity scores. This number significantly exceeds the total number of variants observed in both the general population cohorts and the disease-enriched cohort combined. These scores are predictions of missense variant effects, including substitutions that have not been observed in the datasets used here; they do not establish HCM-specific causality or clinical severity.

#### ClinVar

ClinVar is a public archive that collects and shares information about human genetic variants and their relationships to diseases and drug responses. We used the ClinVar Miner tool to access data on our genes of interest with data current up to February 5, 2024 [20]. In total, this resulted in 5,252 annotations, of which 945 are categorized as pathogenic / likely pathogenic and 1391 are categorized as benign / likely benign. Variants with multiple or discordant submissions were counted as separate annotations. We used data from ClinVar as an independent, high-quality validation set to further substantiate the validity of the identified enriched regions.

### Classification of variants

Our analysis focused on identifying spatially enriched regions using a discrete, binomial statistical framework. We performed an analysis using observed variants and a second analysis supplemented with AlphaMissense predictions. Throughout, disease-observed refers only to variants actually observed in HCM patients.

#### Clinical data: SHaRe vs. population datasets

For the clinical data, we adopted the same categorization strategy as proposed by Homburger et al., labeling all variants found in HCM patients from the SHaRe cohort as disease-observed [1]. Variants exclusively appearing in one of the two population datasets were considered reference variants. While the gnomAD and UKBB population datasets may include individuals with HCM, the incidence is not expected to exceed that of the general population.These labels indicate the source of an observation rather than its clinical classification: a variant observed in an HCM-patient is not necessarily causal, and a population-observed variant is not necessarily benign.

#### Integration of pathogenicity prediction model

The model AlphaMissense provides pathogenicity predictions as a continuous score ranging from 0 to 1. To incorporate these scores into our discrete statistical framework, we ranked predicted substitutions and selected high-scoring and low-scoring subsets. We added these subsets to the HCM-cohort and population-reference classes, respectively, and excluded the remaining predictions. We refer to these combined sets as the augmented HCM and augmented reference classes.

To address the data imbalance between observed variants in clinical datasets and in silico predictions, we implemented a selective inclusion strategy. This approach increased the number of variants included in our analysis by 50%, while maintaining the ratio of HCM-cohort to reference variants constant. Before integrating AlphaMissense data, we performed a deduplication step to remove variants already present in the SHaRe cohort or population datasets. We used a selected subset rather than the full saturation set to limit the contribution of predictions relative to observed variants.

After excluding substitutions already observed in the clinical or population datasets, we ranked the remaining predictions by score. The number of high-scoring substitutions added to the HCM-cohort class was 50% of the observed HCM-cohort variant count; the number of low-scoring substitutions added to the reference class was 50% of the observed population-reference variant count. These additions increased the total number of variants by 50% without changing the class ratio. Selection was based on theirrank predictedand class size rather than a clinical pathogenicity threshold:

*AddedDiseaseVariants = sorted Predictions[0:N_Disease_×0.5]*

Similarly, for population variants, we added the lowest-scoring variants:

*AddedPopulationVariants = sorted Predictions[N_Population_×0.5:N_Population_]*

#### Selection of structural models

We evaluated structures generated using the AlphaFold Protein Structure Database, ColabFold with and without experimental templates, and AlphaFold 3 [14], [21], [22], [23]. Model selection considered predicted local confidence (pLDDT), predicted aligned error (PAE), and alignment with available experimental structures. We selected AlphaFold 2 models for MYH7, MYBPC3, MYL2 and MYL3, an AlphaFold 3 model for TNNT2, and a ColabFold model with templates for TNNI3. MYH7 spatial scanning was restricted to subfragment 1 (S1; residues 1–840). AlphaFold 3 was used for protein-ion and protein-ligand structure predictions. Template identifiers, modeling parameters and gene-specific comparisons are provided in the Supplemental Methods and the supplementary model-evaluation section.

#### Three-dimensional spatial scan

In our binomial spatial scan analysis, we mapped unique variants onto the structural models and compared the proportion in the HCM-cohort class inside each sphere with the proportion outside it. The same procedure was applied to the augmented HCM and reference classes after inclusion of AlphaMissense predictions. Thus, enrichment in the observed data analysis reflects the distribution of patient and population observations, whereas enrichment in the augmented analysis also depends on the selected predictions.

The statistical framework utilizes a sliding-window approach, using spheres with radii of 10Å, 15Å, and 20Å, which iterate across all amino acids of the protein.

For each sphere, we counted HCM-cohort variants within (y_in_) and outside (y_out_) the sphere and population-reference variants within (z_in_) and outside (z_out_). In the augmented analysis, these counts also included the selected high-scoring and low-scoring substitutions, respectively. The proportions were defined as follows:

- p_in_=y_in_/(y_in_+z_in_) : The proportion of variants in the HCM-cohort or augmented HCM class within a given window.
- p_out_=y_out_/(y_out_ + z_out_): The proportion of variants in the HCM-cohort or augmented HCM class outside the window.
- r=(y_in_+y_out_)/(y_in_+z_in_+y_out_+z_out_): The overall rate of variants in the HCM-cohort or augmented HCM class across all windows.

For each window, we compute the binomial log-likelihood ratio statistic. This statistic compares the null model, where p_in_ and p_out_ are both equal to the overall rate r, against an alternative model where p_in_ differs from p_out_. The log-likelihood ratio statistic for each window is calculated as follows:

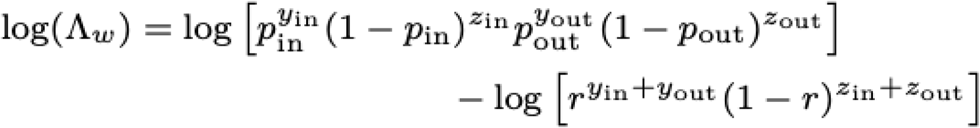

For each residue of the protein model, the likelihood ratio was calculated. We then assessed the statistical significance of the identified enriched regions using a permutation test with 1,000 permutations. During each permutation, the variant labels were shuffled randomly, and the number of variants per residue was held constant. This ensured that the significance assessment remained unaffected by any inherent non-uniform distribution of variants across the full protein.

### Supporting analyses

#### ClinVar validation

We performed a qualitative assessment of ClinVar classifications within and outside the identified regions by examining histograms of the distribution of pathogenic and benign variants across these regions. For quantitative analysis, we compared the proportions of pathogenic and likely pathogenic variants against benign and likely benign variants using Fisher’s exact test.

This comparison assesses agreement with clinical annotations; it does not establish the pathogenicity of each variant in an enriched region.

#### Disease onset age analysis

We compared the age of disease onset in patients with variants located within the identified enriched regions to those with variants outside the sphere. The statistical significance of observed differences was assessed using a Wilcoxon rank-sum test and Kaplan-Meier curves for disease onset.

#### External validation set

We validated our findings with an independent dataset containing variant information of patients diagnosed with HCM. The results of this validation are presented in the supporting materials.

## Results and Discussion

We first quantified the distribution of variants across major sarcomeric genes in both population and patient cohorts, complemented by *in silico* pathogenicity predictions (Table 1). Our framework compares the spatial distribution of variants found in HCM patients to those seen in population cohorts to identify regions of disease enrichment. These regions were then assessed using two complementary approaches: enrichment of pathogenic classifications in ClinVar and differences in age of disease onset in affected individuals. This analysis integrated three large-scale genetic resources: gnomAD (730,947 exomes, with 586,403 individuals carrying variants in the genes of interest), the UK Biobank (500,000 individuals, of whom 346,977 carried variants in these genes), and the SHaRe registry (17,000 patients with cardiomyopathy, including 12,326 with HCM, of whom 4,427 carried variants in the six genes studied). Our analysis revealed strong functional implications of 15 regions enriched for variants across key sarcomeric genes, including the kinetics of myosin ATPase activity, the biophysical properties of the myosin molecule, and the stability of myosin sequestration in the sarcomere (Table 2).

**Table 2.**
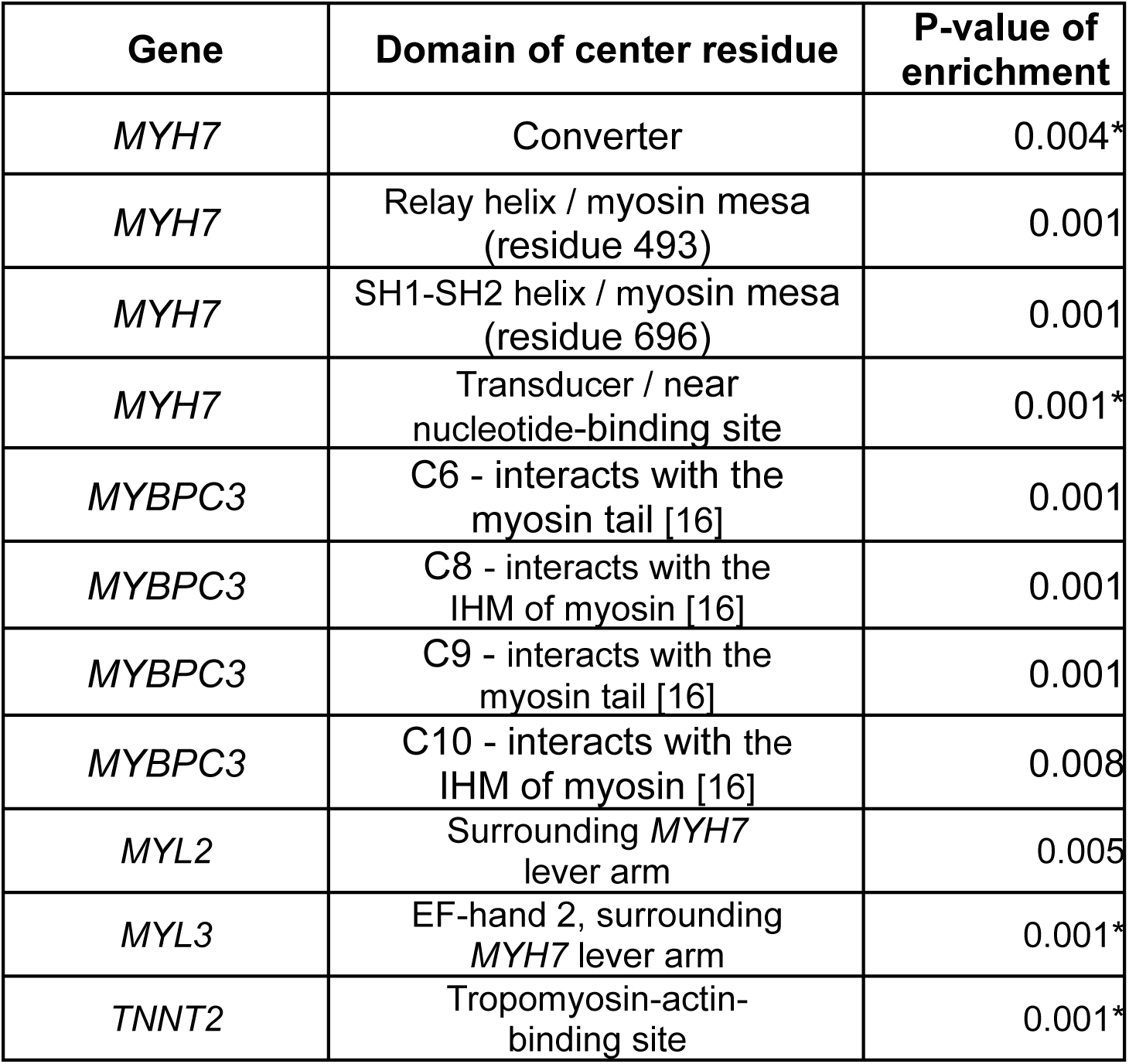
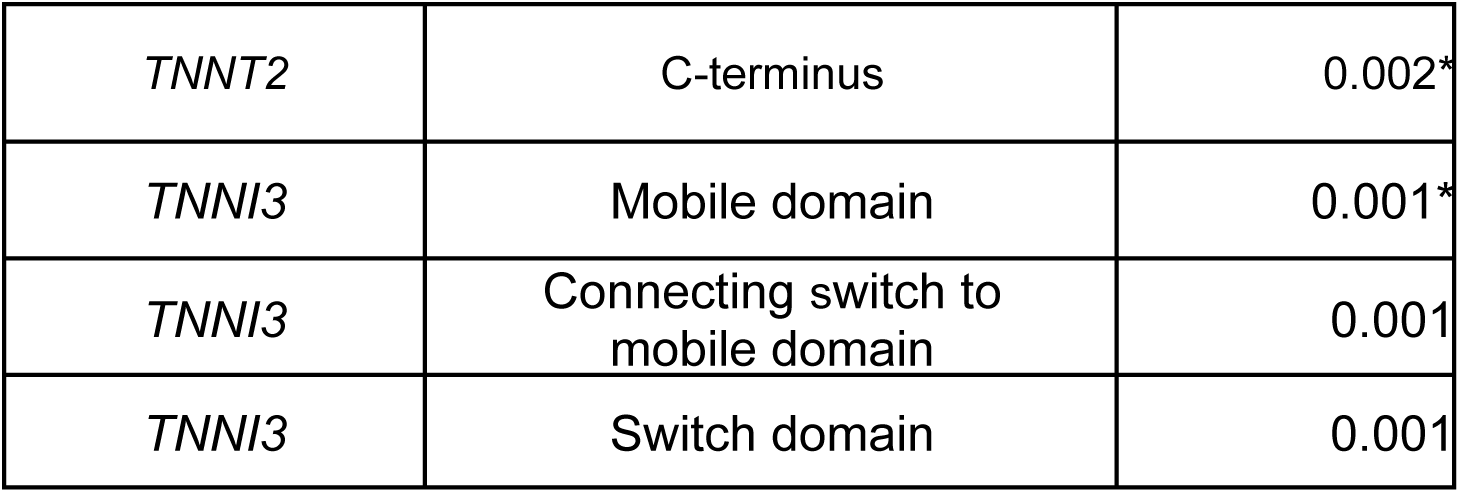
Significance assessments for identified variant enriched regions across different genes and protein domains. For each region, the expected number of variants was estimated under a null model assuming a uniform spatial distribution across the protein structure, and compared to the observed count using permutation testing. P values are two-sided. IHM: Interactive Head Motif *Region was also detectable using real-world clinical and population data only (prior to incorporating pathogenicity prediction scores into the analysis).

### *MYH7* converter and transducer domains are associated with early disease onset

In the Subfragment 1 (S1) domain of the beta myosin heavy chain, we identified two key regions enriched for variants observed in disease (see Fig. 1a). The first region identified was the converter domain, centered on residue 724 with a 15 Å sphere (p=0.004). This region was associated with dramatically earlier disease onset (10.5 years earlier, p=2.8×10−7, see Fig. 2) and showed significant enrichment of pathogenic/likely pathogenic variants in ClinVar (p=0.024).

**Figure 1.**
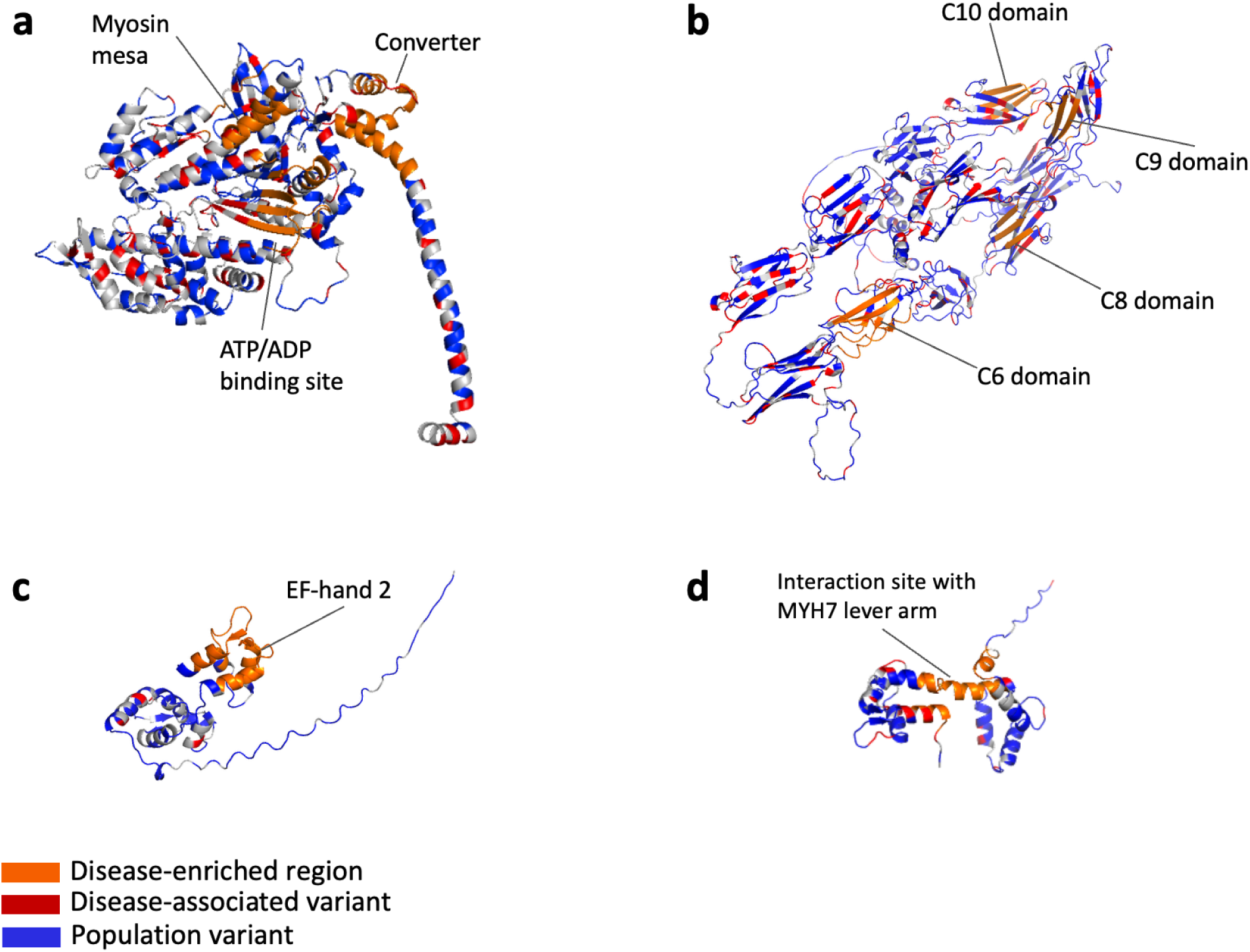
Structural model of MYH7, MYBPC3, MYL2, and MYL3 highlighting regions enriched showing disease-enriched variants in orange, disease-observed variants in red and population variants in blue. (a) Structural model of MYH7 S1 (subfragment 1) region with the regions enriched with disease-observed variants highlighted in orange (myosin mesa, converter, and nucleotide binding site). (b) Structural model of MYBPC3 highlighting disease-enriched regions in the C6, C8, C9, and C10 domains. The augmented analysis identified significant clustering on beta-sheets, supported by a comparison with ClinVar annotations. (c) Structural model of MYL3 highlighting the disease-enriched region on the second EF-hand domain, centered on residue 182. (d) Structural model of MYL2 highlighting the disease-enriched region centered on residue 98 and associated with significantly later disease onset.

**Figure 2.**
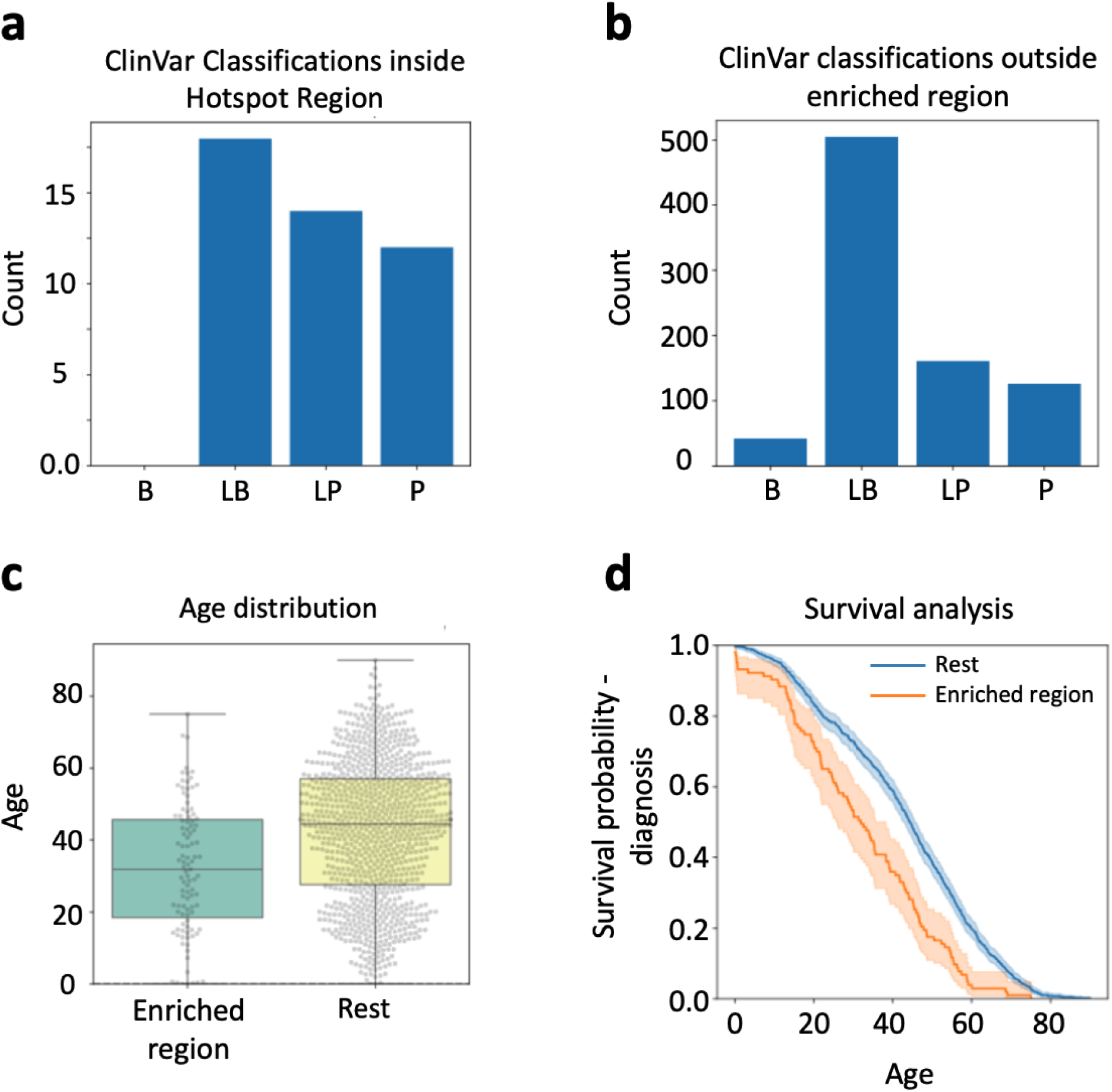
As orthogonal validation of the findings, ClinVar classifications and disease onset analyses demonstrate a significant enrichment of pathogenic/likely pathogenic (P/LP) variants compared to benign/likely benign (B/LB) in the MYH7 converter region, as well as a significantly earlier disease onset (Δ = 10.5 years). (a) Distribution of ClinVar classifications for variants inside the enriched converter region. (b) Distribution of ClinVar classifications for variants outside the converter region. (c) Age distribution at diagnosis for diseased subjects with variants inside vs. outside the converter region. (d) Kaplan–Meier survival analysis showing earlier onset for diseased subjects with variants in the converter region.

The converter domain enrichment aligns with findings from previous studies [1], [24], [25], and our observation of earlier diagnosis is consistent with earlier research [1], [6]. This domain was previously suggested to have a critical role in force transduction during systolic contraction [8] by coupling the biochemical events of ATPase activity and the power stroke, which could explain why variants in this region may cause HCM. However, functional analysis of the recombinant human cardiac myosin with converter domain variants showed surprisingly little change in motor function [26]. Rather, destabilizing the sequestered state of myosin seemed to result in increasing the availability of myosin heads for forming crossbridges [27] which creates the characteristic hypercontractility of HCM.

A second region, previously suggested as enriched but not statistically confirmed, was identified near the nucleotide-binding site centered on residue 257 with a 15 Å sphere (p=0.005). This region is known as the transducer domain, with structural elements at the heart of the motor domain comprising the seven-stranded beta sheet and connectors that control its conformation within the myosin motor [28]. It trended towards an onset of disease 4.2 years earlier onset (p=0.071, see supplemental Figure S7) and showed significant enrichment of pathogenic variants in ClinVar (p=0.004, where pathogenic refers to variants classified as pathogenic or likely pathogenic by ClinVar). Our mathematical characterization significantly strengthens the evidence for a pathogenic region in the ATP/ADP binding region moving beyond prior qualitative descriptions. By quantitatively establishing residue 257 and its surrounding region as a region statistically significantly enriched with disease-observed variants, this analysis deepens our understanding of its clinical importance and functional relevance.

Functional analyses of several variants in the transducer domain have been reported previously. The p.Arg453Cys variant decreases the maximal ATPase activity and reduces actin filament velocity but significantly increases the intrinsic force of the myosin motor domain. Consequently, this variant leads to an overall hypercontractile state in cardiac muscle [29]. Similarly, the p.Arg249Gln variant significantly increases the opening of the interacting heads motif (IHM), releasing sequestered myosin heads, thereby enhancing ATPase activity, as demonstrated by Adhikari et al. [27]. Both variants result in a hypercontractile effect on cardiac myosin function.

### Disease-observed *MYL3* variants cluster on the EF-hand domain

The examination of *MYL3* revealed one region enriched with variants observed in disease on the second EF-hand of the protein, defined by a 15 Å sphere centered on residue 182 (p = 0.001; see Fig. 1c). This region also shows a significantly higher number of pathogenic variants in ClinVar than expected by chance (p = 0.006). Given that this region corresponds structurally to the functional Ca²⁺ binding domain, the enrichment of disease-observed variants suggests significant functional relevance. However, previous studies have indicated that despite the presence of a canonical Ca²⁺ binding motif, *MYL3* has evolutionarily lost its calcium-binding capacity [30], [31], raising uncertainty about the true functional impact of variants in this region. To clarify these conflicting observations, we performed protein-ion interaction simulations to investigate the potential structural and functional implications in greater detail.

Within the enriched sphere, 16 out of the 41 patients observed with a *MYL3* variant carried the p.Glu143Lys variant. This variant was initially reported in a family with early-onset cardiomyopathy with mid-cavitary hypertrophy and restrictive physiology in which siblings carrying the homozygous p.Glu143Lys variant showed severe phenotype [32]. Another patient who was homozygous for *MYL3* p.Glu143Lys and heterozygous for *MYL2* p.Gly57Glu had severe restrictive cardiomyopathy (RCM) [33]. These reported two cases suggest the functional importance of this domain in determining the relaxation of the myocardium that is abnormal in both HCM and RCM. A protein-ion interaction simulation using AlphaFold 3 demonstrated that residue 143 is in close proximity to the defunct divalent cation binding site (see Fig. 3a). Introducing the p.Glu143Lys variant into the amino acid sequence and re-running the simulation showed a displacement of the cation binding site from its original position (see Fig. 3b, Fig. 3c).

**Figure. 3.**
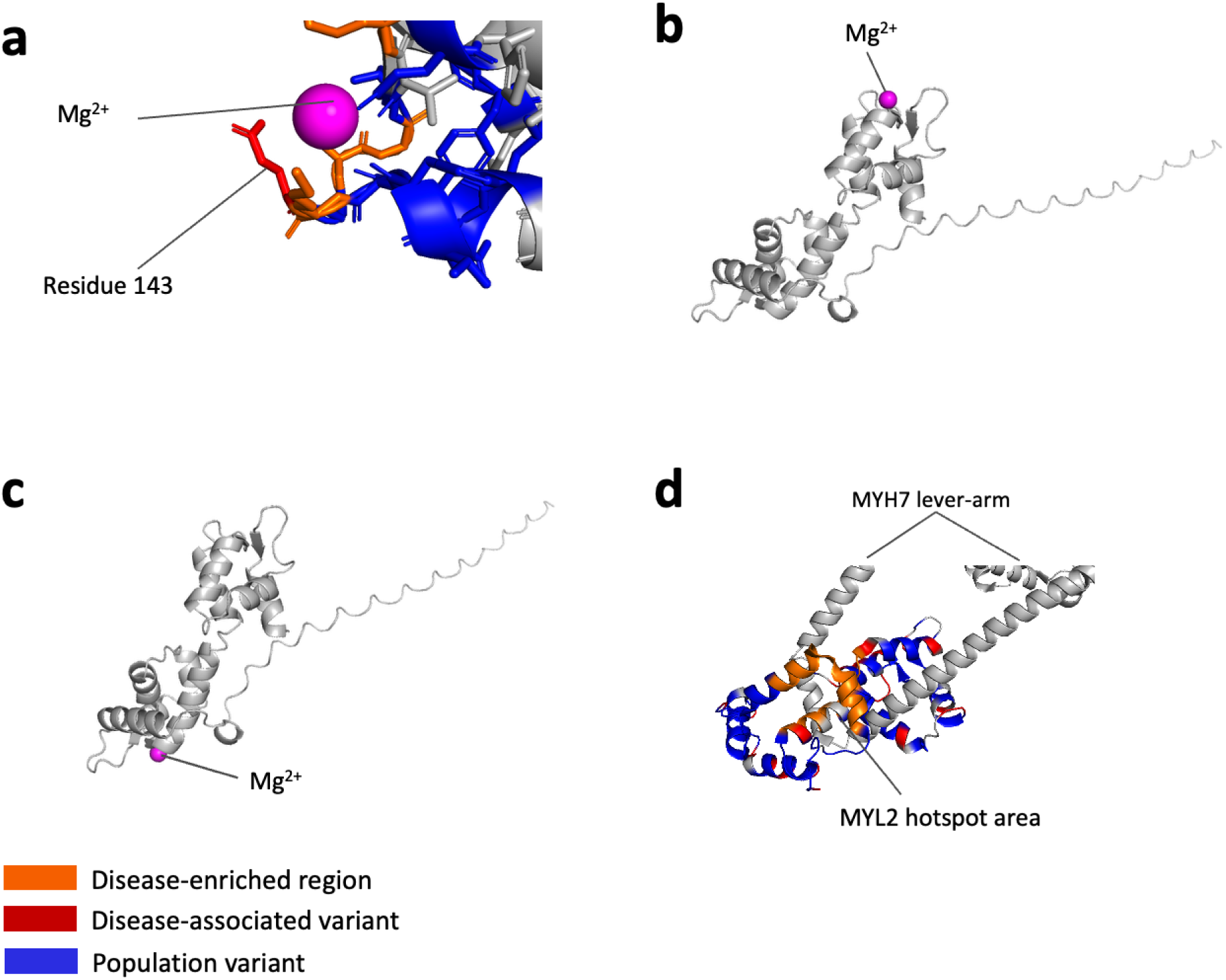
Structural models of MYL2 and MYL3 highlighting the disease-enriched region. (a) Close-up of the potential MYL3 cation interaction site, showing the proximity of residue 143 to the magnesium ion (Mg^2+^). This variant might have its pathogenic effect by disrupting MYL3’s interaction with cation. (b) For the MYL3 wild-type structure, AlphaFold 3 simulates an interaction near residue 143. (c) For the MYL3 structure with variant p.Glu143Lys, the interaction is moved. (d) Close-up of the MYL2 disease-associated region, showing its proximity to the MYH7 lever arm, suggesting that variants in this region may affect the interaction between MYL2 and MYH7 [16].

This evidence raises a hypothesis that substituting the negatively charged glutamic acid with the positively charged lysine influences the affinity of *MYL3* to divalent cations and thereby contributes to functional impairment. In general, EF-hand motifs act as calcium sensors, undergoing conformational changes upon calcium binding, which are essential for cellular function [34]. EF-hand motifs are also able to bind to Mg^2+^, and the introduction of variants can abolish the binding of either cation. The simulated structural changes may suggest that protein-ion interaction is significantly altered by the p.Glu143Lys variant.

Several variants in this region have been shown to affect protein function and stability. The variant p.Glu143Lys has been shown to significantly weaken the binding of *MYL3* to the myosin lever arm [35], and a transgenic mouse model harboring this variant showed increased ATPase activity, duty ratio, and crossbridge formation that led to a significant increase in force generation and diastolic dysfunction [36]. Additionally, the p.Arg154His variant significantly weakens the binding to the myosin lever arm, exhibiting a 3-fold higher KD in protein-binding experiments [35].

Despite human *MYL3* losing its ability to bind Ca²⁺, the enriched region identified on the protein structure matches exactly the evolutionarily lost divalent cation-binding site. Our simulation raises the hypothesis that while *MYL3* does not directly bind Ca²⁺, the evolutionarily conserved second EF-hand still has a regulatory function, possibly by interactions through ions such as Mg²⁺ and a major structural alteration that affects its binding to myosin.

### Integration of variant effect predictions with observed data identifies nine additional enrichment regions

By incorporating in silico pathogenicity predictions (AlphaMissense) into our spatial analysis framework, we expanded our ability to detect variant clustering across protein structures. This integrative approach revealed statistically significantly enriched regions in six sarcomeric genes. Notably, MYL2 and MYBPC3, which showed no clustering using real-world data alone, now exhibited a clear enrichment signal, showing that these regions became detectable after adding selected predictions to the observed data (see Table 1). Computational pathogenicity predictions have been shown to correlate with real-world disease associations and outcomes supporting the validity of integrating such estimates with observed variant data [37], [38]. These additional signals depend on the augmentation strategy and should be interpreted as candidate regions requiring further validation.

### Structural prediction augmentation identified additional clusters on the *MYH7* myosin mesa

Using a combined approach with real-world data and pathogenicity predictions, we identified two additional enriched regions and successfully re-identified the previously observed regions in the converter domain and transducer domain. Notably, the statistical significance for the transducer domain improved, with the p-value decreasing from p=0.005 to p=0.001.

The two additional regions were identified on the myosin mesa (see Fig. 1a). The first sphere was centered on residue 493 with a 10 Å radius (p=0.001) and showed significant enrichment of pathogenic variants in ClinVar (p=0.043). This region falls under the relay helix, a connector essential for allosteric communication between the motor domain and the lever arm [15], [39].

Previous structural modeling and molecular dynamics simulation suggested that the variants at residue 493 (p.Met493Leu) and its surrounding area are expected to cause destabilization of IHM [40]. The second sphere, centered on residue 696 with a 10 Å radius (p=0.001), was associated with 8.4 years later disease onset (p=0.016, see supplemental Figure S6) and also showed significant enrichment of pathogenic variants in ClinVar (p=0.041). The residue 696 resides within the SH1 helix, another critical connector that transmits the structural rearrangements within the motor domain to the converter domain [39]. This region is also predicted to destabilize the IHM, leading to hypercontractility [40].

The myosin mesa findings both confirmed and expanded upon previous research [1], [7]. While earlier studies identified a surface area of interest, our work uncovered a three-dimensional enriched region, providing new insights into the spatial characteristics of this functional zone.

The functional validation, as well as structural modeling and prediction results, provide further support for our approach.

### Four variant clusters revealed in *MYBPC3* near myosin tail and IHM motor domain interfaces

Our analysis of *MYBPC3* revealed four regions of enrichment in the augmented HCM class distributed across different domains of the protein (see Fig. 1b). The first region was identified on the previously reported C6 domain, centered on residue 857 with a 15 Å sphere (p=0.007). Two novel regions were found in the C8 domain, centered on residue 1030 with a 10 Å sphere (p=0.001), and in the C9 domain, centered on residue 1102 with a 10 Å sphere (p=0.001). The fourth region was identified in the previously reported C10 domain, centered on residue 1253 with a 10 Å sphere (p=0.008). Although all four regions showed a higher-than-expected proportion of pathogenic variant annotations, only the C6 domain reached statistical significance (p=0.003), and none of the regions were associated with a significant difference in disease onset.

These regions did not show statistically significant enrichment when only mapping real-world data to the 3D protein structure. This lack of significance could be due to the variability in disease severity and later onset associated with *MYBPC3* variants, and underscores the enhanced sensitivity achieved by incorporating *in silico* pathogenicity predictions into our spatial analysis.

The enrichments identified in the augmented analysis in domains C6-C10 are particularly interesting due to their established interaction with the light meromyosin (LMM) region of myosin that forms the myosin tail in thick filament, which is critical for maintaining the stability of the sequestered (inactive) state of myosin (Fig. 4a) [16]. Dutta et al. reported the cryo-EM structure of the human cardiac thick filament in a relaxed state, which revealed the packing of myosin tails in the filament backbone and their interaction with *MYBPC* and titin [16]. The fitted atomic model showed interactions between domains C6-C10 and myosin tail. In particular, domains C8 and C10 interacted with the motor domain of IHM myosin, suggesting that these two domains are involved in the stabilization of IHM. While the C6 and C9 domains do not directly interact with the IHM myosin itself, their interactions with the LMM region may affect the structure of myosin tails and affect stability of myosin head sequestration. Taken together, these results suggest that *MYBPC3* C6–C10 variants may alter sarcomeric structure in a manner that increases myosin head availability, thereby contributing to hypercontractility in HCM.

**Figure 4.**
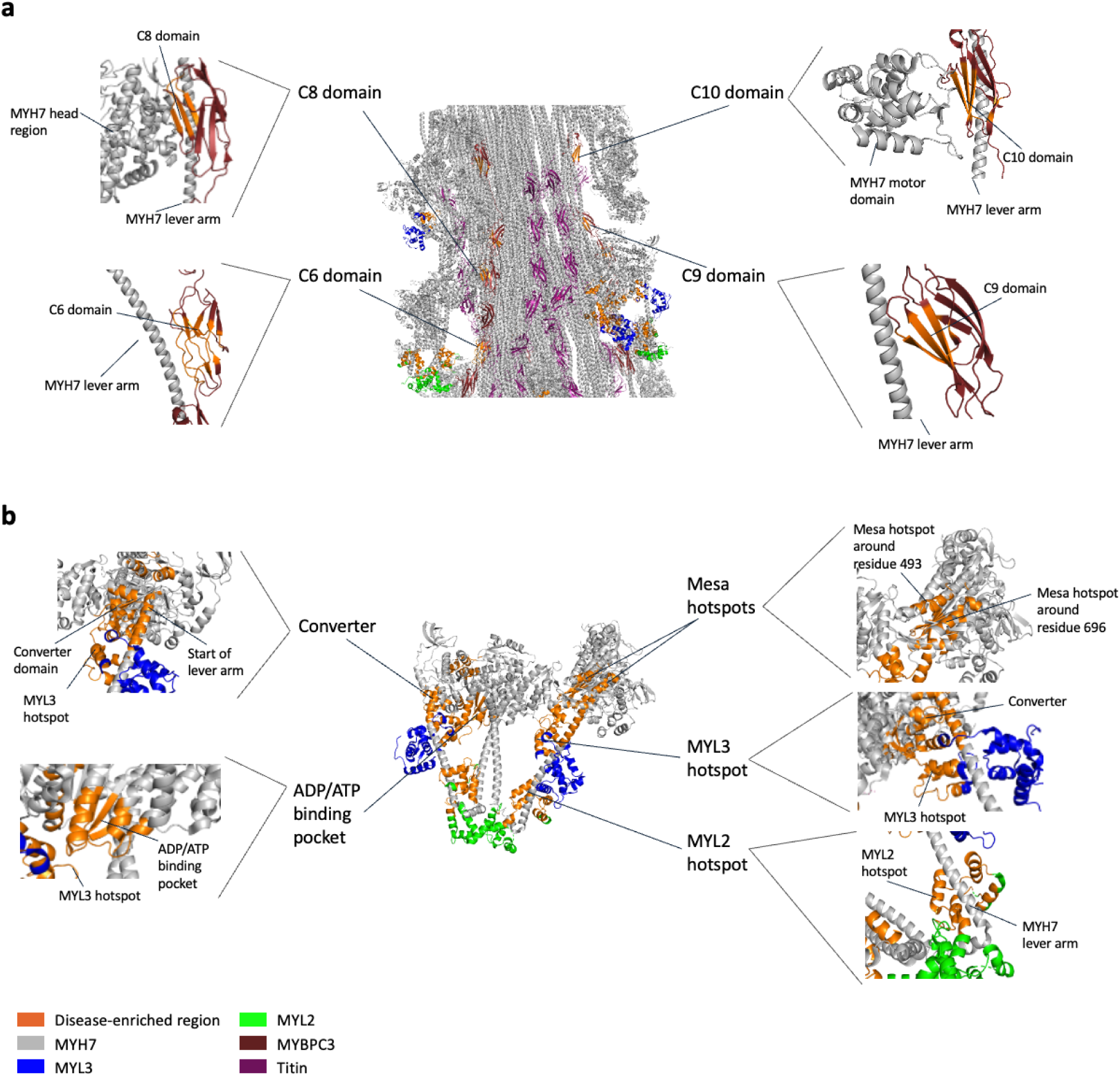
Cryo-EM structures place disease-enriched regions within the human cardiac thick filament structure (a) Cryogenic electron microscopy (Cryo-EM) structure of the human cardiac myosin filament with zoomed in visualizations of the regions identified on MYBPC3 [16]. The four disease-associated regions located in domains C6 (residue 857), C8 (residue 1030), C9 (residue 1102), and C10 (residue 1253) are highlighted, illustrating their close proximity to the myosin tails within the thick filament backbone. (b) Cryo-EM structure of the human beta-cardiac myosin folded-back off state displaying the regions identified on MYH7, MYL2 and MYL3 [15]. The highlighted regions indicate proximity to critical structural elements, including the converter, myosin mesa and transducer domains in MYH7, the lever-arm binding interface in MYL2, and the divalent cation-binding site in MYL3, underscoring their potential functional impact on myosin motor regulation.

Upon examining the disease enriched regions illustrated in Fig. 1b and Fig. 4a, it is also apparent that these regions are predominantly localized within regions where the secondary protein structure consists of beta-sheets. A t-test supports this observation, indicating that the clustering of variants observed in the augmented HCM class on beta-sheets is highly unlikely to occur by chance (p=1×10^−6^). This result is further supported by comparing the pathogenic and benign/likely benign (B/LB) annotations from ClinVar (p=0.049), providing a complementary comparison with clinical variant annotations (see Fig. 1b).

Previous studies analyzing disease regions on *MYBPC3* using two-dimensional sequence-based analyses rather than three-dimensional structural mapping have shown varying results. Several studies specifically compared HCM cohorts to gnomAD as a reference population. Waring et al. identified clusters in the C1, C3, C7, and C10 domains [11], while Helms et al. found variant concentration in the C3, C6, and C10 domains [2]. Walsh et al. observed clustering restricted to the C3 and C10 domains [41]. The C10 domain, in particular, has been consistently highlighted as a potential region for disease-associated variants across multiple studies [2], [11], [41]. However, the significance of other regions, such as C6, C8, and C9, remains less certain. Notably, some studies have reported no significant clustering patterns in any domain [42], [43]. These discrepancies may stem from differences in sample sizes, statistical methods, or the later-onset and variable penetrance of *MYBPC3* variants, which are often truncating and associated with a more favorable prognosis [44].

Our discovery that both the identified disease-associated regions and individual variants classified as pathogenic in ClinVar are disproportionately located on beta-sheets represents a valuable insight into the structural basis of variant pathogenicity in *MYBPC3* (see Fig. 1b). One possible explanation for this clustering on beta-sheets could be the structural role they play in protein stability. Helms et al. identified several *MYBPC3* missense variants that markedly decreased protein stability, suggesting that haploinsufficiency due to rapid degradation could underlie their pathogenicity [2]. Beta-sheets often form the core of protein structures, and disruptions in these regions may more readily lead to misfolding or destabilization, thereby triggering pathogenic effects [45], [46]. Another hypothesis is that beta-sheets may be involved in important protein-protein interactions, such as the ones suggested from the thick filament structure described above, and variants in these regions could disrupt essential functional interfaces between myosin. Further research is needed to investigate whether the clustering in beta-sheets of *MYBPC3* could be linked to disruptions in its structural integrity or functional binding interfaces.

### Variant enrichment in *MYL2* surrounding the *MYH7* lever arm

With integration of a pathogenicity prediction model, our analysis revealed a novel disease-region enriched in the augmented HCM class characterized by a 20 Å sphere centered on residue 98 (p = 0.005), associated with 12.8 years later disease onset (p = 0.005, see Fig. 1d, see supplemental Figure S12). This region is in close proximity to the *MYH7* lever arm (see Fig. 3d), and encapsulates the phosphorylation site on residue Serine 15 [47].

Previous studies have hypothesized that variants within *MYL2* significantly influence cardiac contractility by altering interactions with the myosin lever arm, potentially modifying its mechanical stiffness [48], [49]. Our findings, identifying a region closely surrounding the *MYH7* lever arm, support this stiffness hypothesis, suggesting that variants within this region may similarly impact lever-arm stiffness and consequently influence mechanochemical properties. These results underscore the critical role of the *MYL2*–*MYH7* interaction and highlight opportunities to further investigate how increased mechanical stiffness may enhance the intrinsic force of individual myosin molecules, ultimately contributing to greater ensemble sarcomere force in the pathogenesis of HCM.

### Methodological considerations and future directions

In this study, we conducted the largest-scale analysis of genetic data on HCM to date, encompassing over 814,392 population-based genome sequences alongside more than 10,000 targeted exome sequences from HCM patients. For the first time, we integrated pathogenicity predictions from a transformer-based pathogenicity prediction model (AlphaMissense) into the analysis, expanding on traditional approaches that use clinical and population data and functional validation, and introducing a framework to interpret predictive data in a structural context. While the prediction model does not distinguish between HCM and Dilated cardiomyopathy (DCM), our downstream phenotype analyses focused on HCM provides disease-specific context for the findings.

### Limitations of predicted pathogenicity and clinical severity

AlphaMissense scores estimate missense variant pathogenicity in a general setting [13]. They do not specify the molecular mechanism or distinguish gain of function, loss of function, dominant-negative effects or haploinsufficiency. Different alterations in the same sarcomeric gene can produce different phenotypes, including HCM and DCM. A high score therefore does not establish that a substitution causes HCM or explain how it affects sarcomere function.

Conversely, a low-scoring substitution is not proven benign. The score and the analytical class assigned here should be distinguished from a clinical classification of pathogenic or likely pathogenic.

The augmentation procedure adds predictions rather than independent patient observations. The selected fraction and rank-based inclusion rules may influence the regions identified, and converting continuous scores into two classes discards information about intermediate predictions. Regions detected only after augmentation should therefore be interpreted as candidates for further investigation.

Future validation strategies could include multiplexed assays of variant effect [50], particularly when combined with assessment in human-induced pluripotent stem cells (hiPSCs). These assays would allow many substitutions within the identified regions to be tested experimentally, including substitutions that have not been observed in patients. The measurement of relevant biomarkers, such as B-type natriuretic peptide (BNP) levels [51], could further support these findings by linking molecular alterations to functional cardiac phenotypes. BNP is released in response to increased wall stress and myocardial strain, thus monitoring BNP levels in variant carriers or hiPSC-derived cardiomyocytes could provide a functional readout of contractile imbalance or impaired relaxation. These approaches would not only help confirm the identified regions but also contribute to our understanding of disease pathways in HCM.

Our analysis identified several new disease enrichments and confirmed previously reported ones. Additionally, by examining cryo-EM structures and performing simulations with AlphaFold 3, we obtained further structural evidence supporting the proposed hypotheses on the functional impacts and disease mechanisms of these enriched regions. Moreover, comparisons with ClinVar annotations provided supporting evidence for the identified regions. Our approach highlights the potential utility of in silico models in prioritizing regions for genetic variant investigation and underscores the importance of integrating the data with structural models of the proteins for more comprehensive disease association studies. The identification of new regions and detailed structural insights into their functional consequences could inform the development of novel therapeutic targets for treating hypertrophic cardiomyopathy. Further validation is needed to establish whether these regions can contribute to variant classification within the American College of Medical Genetics and Genomics (ACMG) guidelines and ClinGen framework [52], [53].

## Supporting information

Supporting information on methodology

## Non-standard Abbreviations and Acronyms

DCM: Dilated cardiomyopathy
HCM: Hypertrophic cardiomyopathy
RCM: Restrictive cardiomyopathy
IHM: Interacting heads motif
LMM: Light meromyosin
HMM: Heavy meromyosin
MYBPC: Myosin-binding protein
C MAF: Minor allele frequency
UKBB: UK Biobank
RMSE: Root mean square error
PAE: Predicted aligned error

## Acknowledgments

We thank the investigators and participants in the SHaRe for their support of this project.

## Supplemental Material

Supplemental Results and Discussion: TNNT2 and TNNI3 Experiments

Figures S1–S18

Table S1

## Data and code availability

The clinical genetic data used in this study are derived from the Sarcomeric Human Cardiomyopathy Registry (SHaRe), UK Biobank, and gnomAD. Due to patient privacy and ethical restrictions, individual-level clinical data cannot be made publicly available.

Access to SHaRe data is subject to approval by the SHaRe governance committee. UK Biobank data are available through application to the UK Biobank (https://www.ukbiobank.ac.uk). gnomAD data are publicly available at https://gnomad.broadinstitute.org. ClinVar data were accessed via ClinVar Miner and are publicly available.

All custom code used for data processing, spatial scan statistics, structural mapping, and figure generation in this study is publicly available at GitHub: https://github.com/ChristianCadisch/Spatial_genomics/. The repository contains the full analysis pipeline required to reproduce the computational framework described in the manuscript.

## Author contributions

C.M.C. developed and implemented the computational framework, performed the analyses, generated the figures, and wrote the manuscript. V.P., A.S., M.K., and E.A.A., contributed to study design, interpretation of results, and manuscript revision. All remaining authors contributed clinical data through the SHaRe registry and provided domain expertise and feedback. All authors reviewed and approved the final manuscript.

