## Supporting information on methodology for "Spatial genomics of the cardiac sarcomere"

### Supporting materials

#### Development of protein models

##### *Computational modelling approach*

To obtain optimal structural models for each protein under study, we evaluated four computational modeling approaches and selected the most accurate models based on a set of evaluation criteria.

We first obtained pre-computed models from the AlphaFold Protein Structure Database, a collaborative effort by Google DeepMind and European Bioinformatics Institute (EMBL-EBI) [1]. These models were generated using the AlphaFold Monomer v2.0 pipeline and last updated on November 1, 2022 [2]. Secondly, we utilized ColabFold v1.5.5 to create structure predictions based on the AlphaFold2 v2.2 model weights and using the faster homology search of MMseqs2 [3].

We attempted to enhance the prediction accuracy of the ColabFold model by providing experimentally obtained template structures. We utilized Stanford's shared computing cluster, Sherlock, for GPU access. ColabFold was set to use a custom template mode with MMseqs2 browsing UniRef and environmental datasets. The number of recycles was set to 3 with early stopping tolerance on auto, and relax was limited to 200 iterations. We employed a greedy pairing strategy, and dropout was disabled.

Lastly, we used the AlphaFold Server to create structure predictions with AlphaFold 3 [4]. For all protein-ion and protein-ligand interactions, we used AlphaFold 3. For the analysis of *MYH7*, we focused exclusively on subfragment 1 (S1), encompassing residues 1-840, as this region possesses distinct functional significance and is highly enriched with variants observed in disease.

##### *Protein template selection*

For modeling the beta-myosin heavy chain (encoded by *MYH7*), we provided the model with template structures with PDB IDs 8ACT [5], 8EFH [6], 8EFI [7], 8ENC [7], and 8G4L [8]. For modeling the myosin-binding protein C (encoded by *MYBPC3*), we used template structures with PDB IDs 6CXJ [9], 6G2T [9], 7LRG [10], 7TIJ [11], and 8G4L [8]. For the essential light chain (encoded by *MYL3*) and regulatory light chain (encoded by *MYL2*), we used template structures with PDB IDs 5TBY [12], 8ACT [5], and 8G4L [8]. For modeling the cardiac troponin T (encoded by *TNNT2*) and troponin I (encoded by *TNNI3*), we used template structures with PDB IDs 6KN7 [13], 6KN8 [13], 7UTI [14], and 7UTL [14].

##### *Assessment and comparison of predicted protein structures*

To assess and compare the quality of all protein models, we analyzed predictive and experimental metrics. The structure prediction models provided two predictive scoring methods: predicted local distance difference test (pLDDT), giving a per-residue measure of local confidence, and the predicted aligned error (PAE), giving an estimate of the expected error between each pair of amino acids. We calculated the mean pLDDT and mean PAE for each protein model to understand overall accuracy and local confidence levels. Furthermore, we aligned the predicted structures with partial experimentally determined protein structures available in PDB and measured the minimal root mean square error (RMSE) between the predicted and experimental structures. Visualizations and alignments were made using PyMol 2.5.8.

#### Result evaluation and selection of 3D protein models

##### *MYH7*

The AlphaFold 2 structure was selected as the best structural model for *MYH7*, demonstrating superior performance across multiple evaluation metrics. With a mean pLDDT score of 73.75 and mean PAE of 23.43 Å, the AlphaFold 2 model achieved the lowest overall PAE and the second-highest pLDDT among all evaluated models (detailed PAE plots and RMSE comparisons available in the supporting information).

All models consistently showed higher accuracy in predicting the structure of subfragment 1 (S1) of the heavy meromyosin (HMM) domain, which encompasses the myosin head (residues 1-840). In contrast, the subfragment 2 (S2) (residues 841-1280) and the light meromyosin (LMM) domain (residues 1281-1935) exhibited lower confidence, with higher predicted errors in these regions.

When aligned with experimentally determined cryo-EM structures, the AlphaFold 2 model consistently produced the lowest RMSE across the tested structures. The model achieved an RMSE of 1.35 Å for the structure with PDB ID 8ACT [5], 3.73 Å for PDB ID 8ENC [7], 3.71 Å for PDB ID 8EFI [7], 6.39 Å for PDB ID 8G4L [8], and 3.25 Å for PDB ID 8EFH [6]. With an average RMSE of 3.67 Å, this model achieved the lowest overall error across the evaluated structures.

##### *MYBPC3*

In modeling *MYBPC3*, all models exhibited similar mean PAE scores, indicating comparable overall accuracy in predicting residue-residue distances across the protein. Additionally, all models, except for ColabFold v2.3 with templates, achieved similar pLDDT scores close to 80, demonstrating high confidence in local structural predictions. The AlphaFold 2 structure, with a mean pLDDT of 79.38 and a mean PAE of 25.43, was ultimately chosen as the best model due to its consistently good performance in terms of RMSE when aligned with experimentally determined cryo-EM structures (detailed PAE plots and RMSE comparisons available in the supporting information).

A detailed inspection of the PAE plots across all models revealed that while the confidence in long-distance relationships between residues far apart in the sequence is low, the local accuracy for individual domains of *MYBPC3* is high. This pattern suggests that, although the models may struggle with accurately predicting the overall topology and domain-domain interactions, they reliably capture the secondary structures and the shape of the individual domains.

The structural differences observed between our *MYBPC3* model and that reported by Thompson et al. likely arise from distinct computational modeling approaches employed [15]. Specifically, Thompson utilized the STRUM algorithm combined with I-TASSER structural predictions focused on individual protein subdomains, potentially highlighting localized folding instability, whereas our integrative approach emphasized whole-protein AlphaFold predictions and spatial clustering analyses across larger protein domains.

##### *MYL2*

For the gene *MYL2*, we selected the model created with AlphaFold 2 as the best structure. It achieved the best mean pLDDT of 83.36 and the second-best mean PAE with 13.21 Å, only surpassed by the structure created with AlphaFold 3 which achieved a mean PAE of 11.49 Å. The decision to use the AlphaFold 2 model was driven by the excellent RMSE when the structure was aligned to the cryo-EM structure with PDB ID 8ACT [5], which itself has a high resolution of 3.6 Å. An inspection of the PAE plots reveals that all structure prediction models are confident in the aligned distances in the domain of the EF-hands: the first EF-hand between residue 24 to 59, and the second and third EF-hands between residues 94-129 and 130-165.

##### *MYL3*

Upon evaluating the structural predictions of *MYL3*, the AlphaFold 2 model was identified as the most reliable, achieving a high mean pLDDT of 88.86 and a mean PAE of 11.14 Å, which, while third lowest, is comparable to the lowest PAE of 10.63Å. This model exhibited exceptional alignment with the cryo-EM structures used for comparison. When aligned with the structure having PDB ID 8ACT [5], the model achieved an RMSE of 1.48 Å, and with the structure having PDB ID 5TBY [12], it achieved an RMSE of 1.12 Å. The PAE plots indicate that, apart from the disordered N-terminal region (amino acids 1-37), the predicted folding confidence is very high.

#### Spatial scanning of 3D protein structure integrated with clinical and population data reveals variant clusters in *TNNT2* and *TNNI3*

##### *Disease-associated TNNT2 variants cluster on tropomyosin-actin interaction site and C-terminus*

For cardiac troponin T, we identified two regions enriched with disease-associated variants (see Figure S1). The first region is located at the edge of the domain where TNNT2 interacts with tropomyosin and actin, defined by a 15 Å sphere centered on residue 95 ($p = 0.002$). The interaction of TNNT2 with actin based on the cryo-EM structure from PDB ID 7UTL [14] is shown in Figure S1(b). The proximity of the enriched region, colored in orange, suggests variants in this location might disturb the interaction. Variants in this region are associated with an earlier disease onset, averaging 10.2 years earlier ($p = 0.003$). This finding is further supported by a higher proportion of P/LP variant annotations in ClinVar ($p = 0.001$).

The second region is situated at the C-terminus of the gene, with a 15 Å sphere centered on residue 290 ($p = 0.003$). Variants in this region are linked to a later disease onset, averaging 8.5 years later ($p = 0.005$), with no significant increase in the proportion of P/LP variant annotations in ClinVar.

##### *TNNI3 variant cluster with earlier disease onset identified on mobile domain*

Upon examining cardiac troponin I, we identified a significant variant enrichment in the mobile domain ($p = 0.001$, see Figure S2). Variants in this region were associated with disease onset 9.6 years earlier ($p = 0.006$) and showed a significantly higher proportion of P/LP variant annotations in ClinVar ($p = 0.001$). In vitro motility assay studies have shown that the deletion of the last 17 residues was associated with myocardial stunning and increased $Ca^{2+}$ sensitivity [16].

Incorporating data from an *in silico* pathogenicity model (AlphaMissense [17]) identified two additional regions enriched in the augmented HCM class on *TNNI3* and also identified the mobile-domain enrichment (see Figure S2). The first additional variant cluster was located within the switch domain, defined by a 10 Å sphere centered on residue 141 ($p = 0.001$). Molecular dynamics simulations found that the p.Arg145Trp variant, responsible for 48% of patients within the enriched area, led to a reduction in interaction between this residue and cardiac troponin-C [18]. The electron microscopy image derived from the PDB with ID 7UTI

[14] also shows the proximity of this enriched region to the actin filament. Variants in this region might therefore also disturb the interaction of *TNNI3* with actin.

The second additional enriched region was defined by a 10 Å sphere centered on residue 174, where the switch domain connects to the mobile domain ($p = 0.001$). This region has been shown to be essential for the interaction with actin and stabilizing the off-state of regulatory units at low levels of $Ca^{2+}$ [19].

#### Validation with external dataset

In addition to validation using ClinVar, we sought further evidence to corroborate our findings by leveraging an independent dataset. We collaborated with the group of Prof. James Ware to get an additional dataset containing variant information from HCM patients, providing an external dataset for comparison. To assess the robustness of our cluster identification, we tested the hypothesis that within identified enriched regions, the observed number of variants exceeds the expected number of variants under a null model assuming a uniform distribution across the protein structure. This analysis confirmed 11 out of 15 previously identified regions (Table S1), reinforcing our findings. In the four regions where the finding was not corroborated, the expected number of variants was low (≤2), which likely limited the power of the validation, as such small sample sizes introduce variability that may skew the results.

#### Evaluation and selection of 3D protein models

##### *TNNT2*

In the evaluation of structural models for *TNNT2*, all models showed comparable PAE scores around 22 Å. The pLDDT scores for the models predicted using the AlphaFold 2 architecture (79.75, 78.24, and 78.22) were higher compared to the AlphaFold 3 model, which achieved a pLDDT of 73.32. Yet, the AlphaFold 3 structure was ultimately selected as the best model due to its superior performance in terms of RMSE when aligned with the experimentally determined cryo-EM structure. It achieved an RMSE of 2.49 Å for the structure with PDB ID 7UTI [14], beating the second-closest structure created with ColabFold v2.3 by 0.5 Å. The PAE plots across all models highlight that the models reliably capture the folding and structural integrity of the three coil segments (residues 1-50, residue 100-180, and residues 220-280) while displaying a lower confidence in predicting the relationships between these domains.

##### *TNNI3*

The ColabFold v2.3 model with templates achieves the best evaluation criteria in the structural modeling of *TNNI3*. The model achieved a pLDDT score of 81.19 and a PAE of 17.6, both of which are superior to those of all other predicted structures. The outstanding performance is also reflected in achieving the lowest RMSE when aligned with the experimentally determined cryo-EM structure, achieving an RMSE of 0.91 when aligned to the structure with PDB ID 4Y99 [20]. An inspection of the PAE plot reveals an especially high confidence of residue-residue distances between amino acids 20 and 150, where both the TNC-binding and actin-binding sites are.

#### Figures

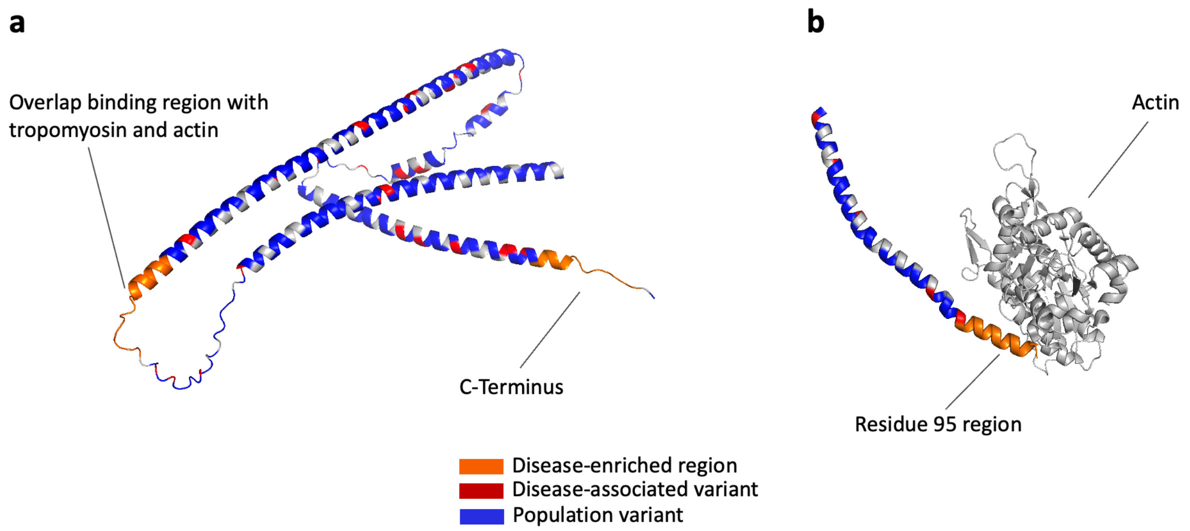


***Figure S1**. AlphaFold 2 structural model of TNNT2 highlighting disease-enriched regions. (a) The C-terminus and overlap binding region with tropomyosin and actin show regions of disease-associated variants. (b) Close-up of the residue 95 region, where TNNT2 interacts with actin [14].*

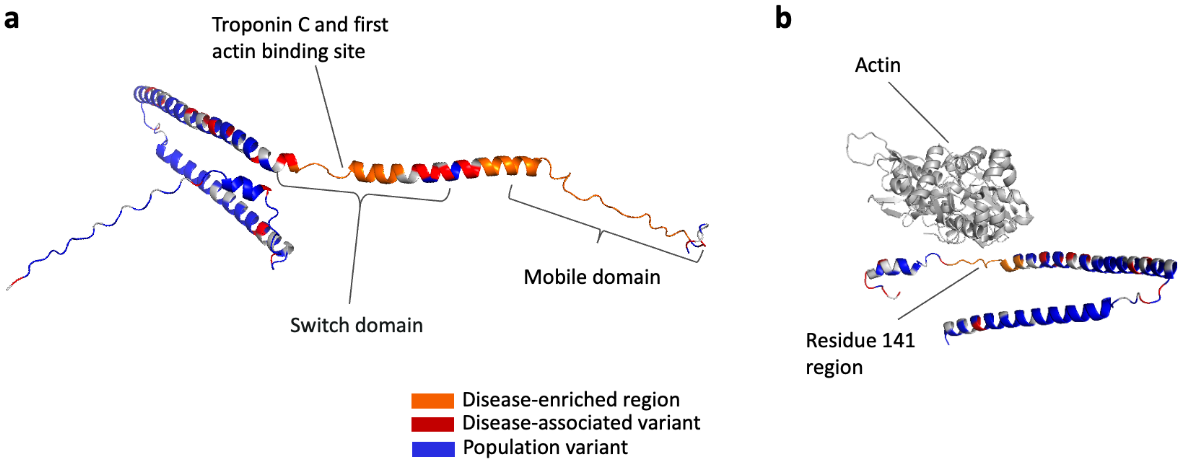


***Figure S2***. *AlphaFold 2 structural model of TNNI3 highlighting disease-enriched regions. (a) Three regions are identified across adjacent domains: the switch domain, the mobile domain, and the Troponin C/first actin binding site. (b) Close-up of the residue 141 enriched region, where TNNI3 interacts with actin [14].*

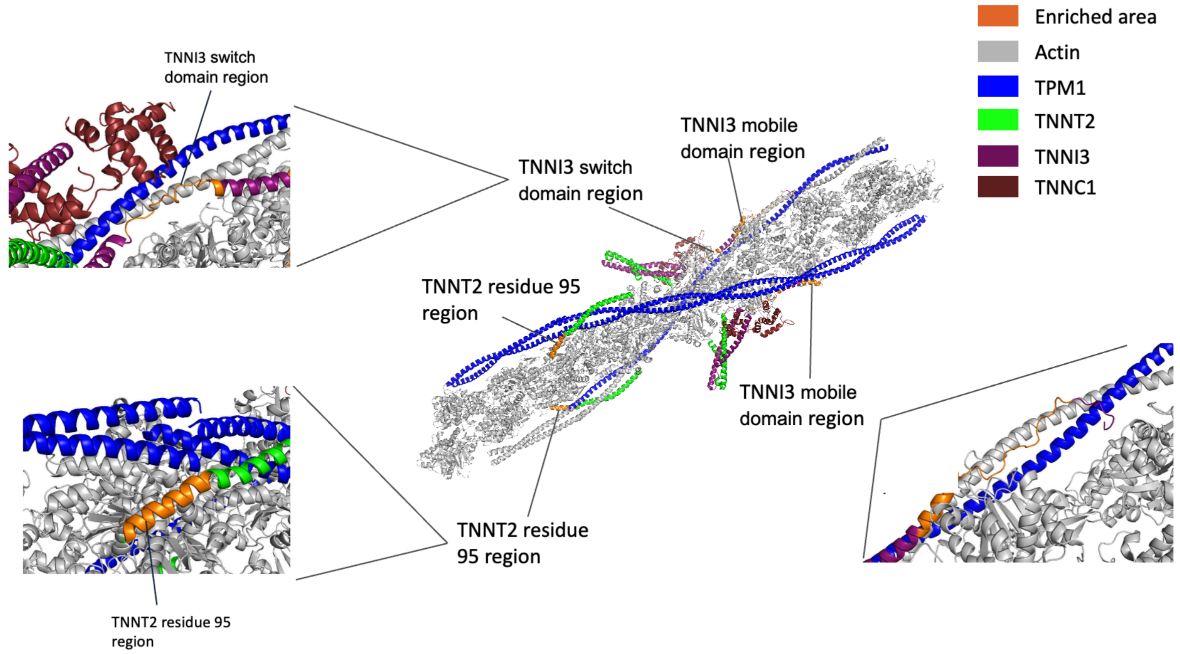


***Figure S3***. *Cryo-EM structure of tropomyosin in human cardiac thin filament in the calcium-free state with the enriched regions identified for TNNI3 and TNNT2 highlighted [21]*

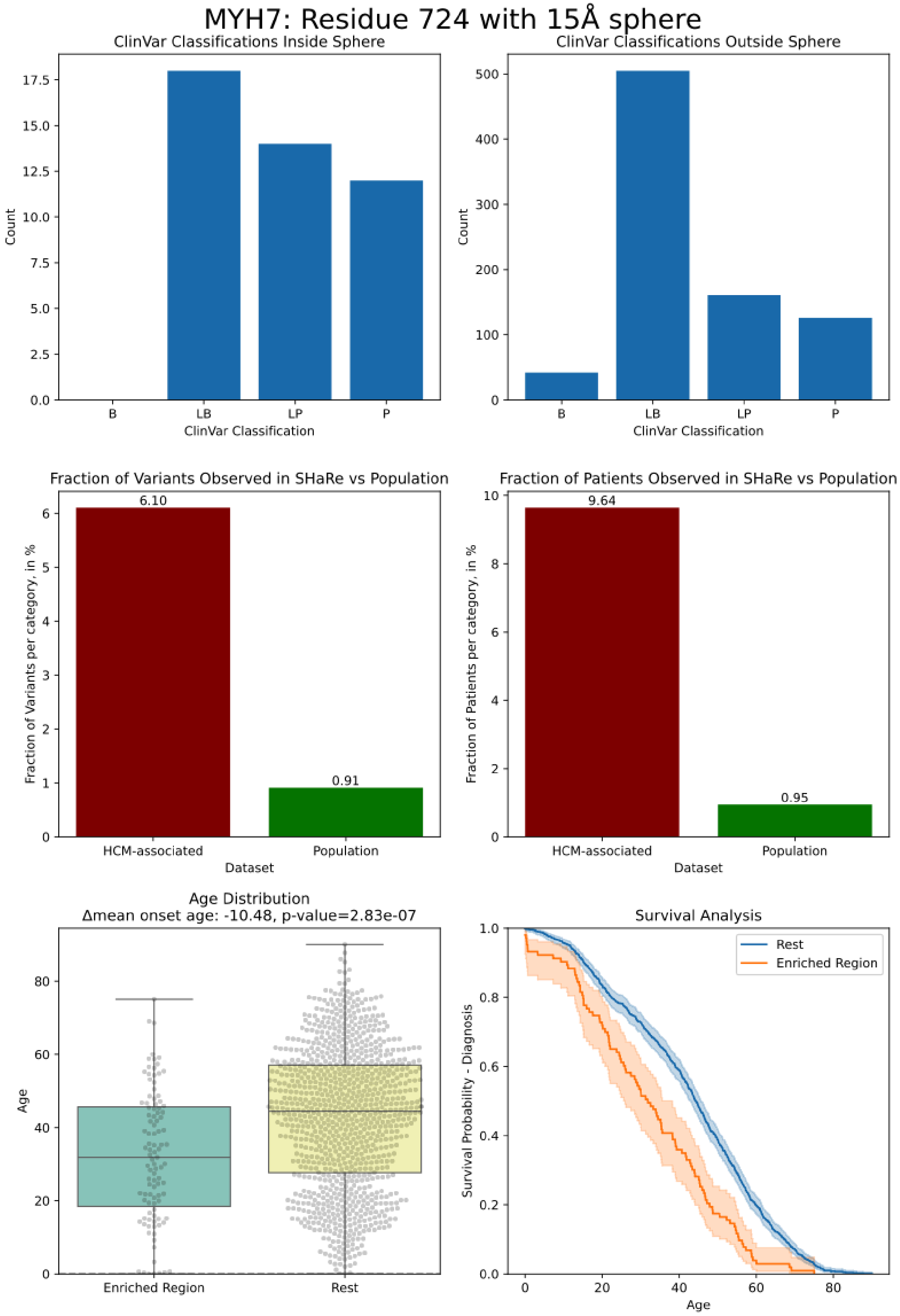


***Figure S4.*** *Validation plots for MYH7 converter domain cluster located on residue 724*

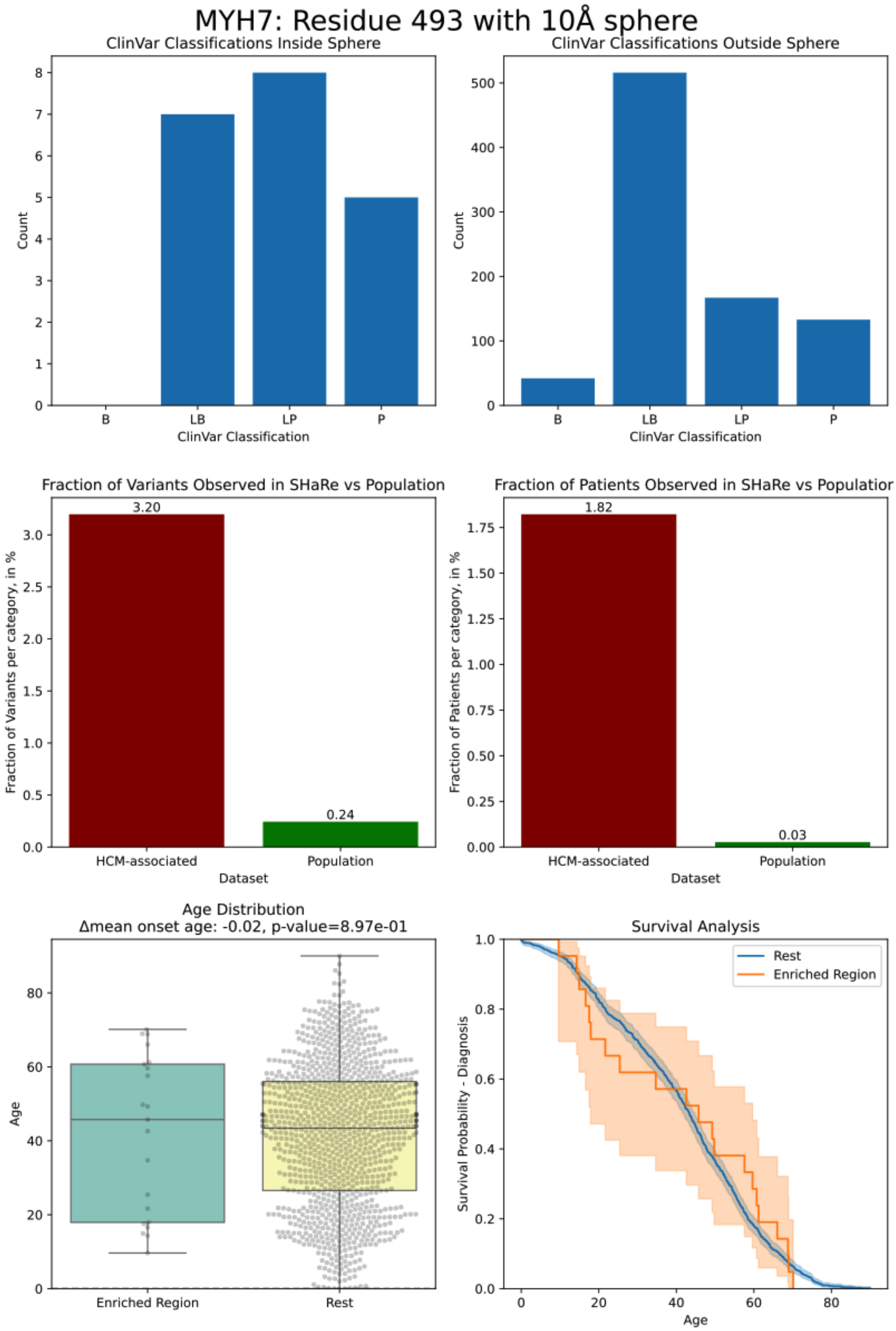


***Figure S5.*** *Validation plots for MYH7 myosin mesa cluster located on residue 493*

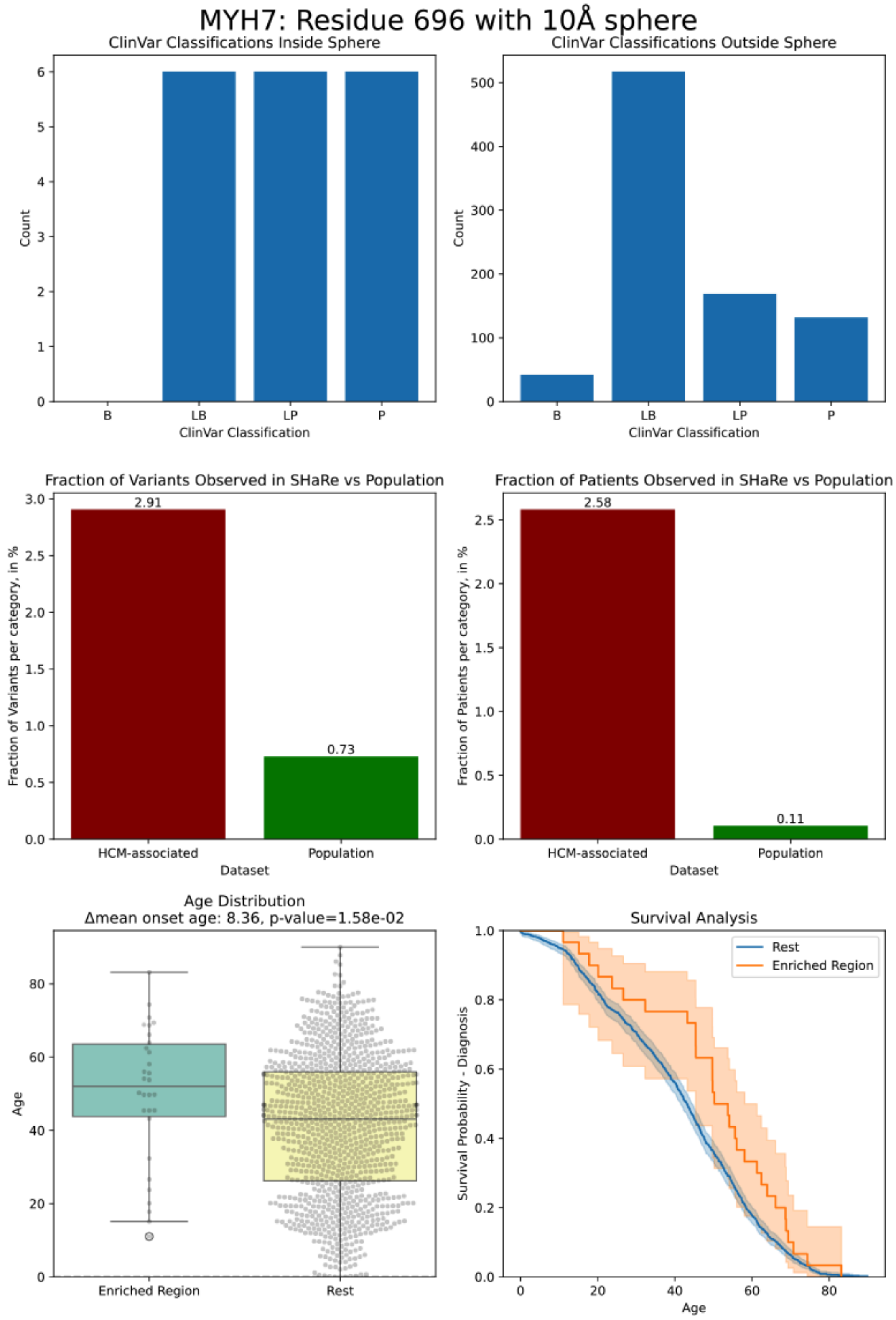


***Figure S6.*** *Validation plots for MYH7 myosin mesa cluster located on residue 696*

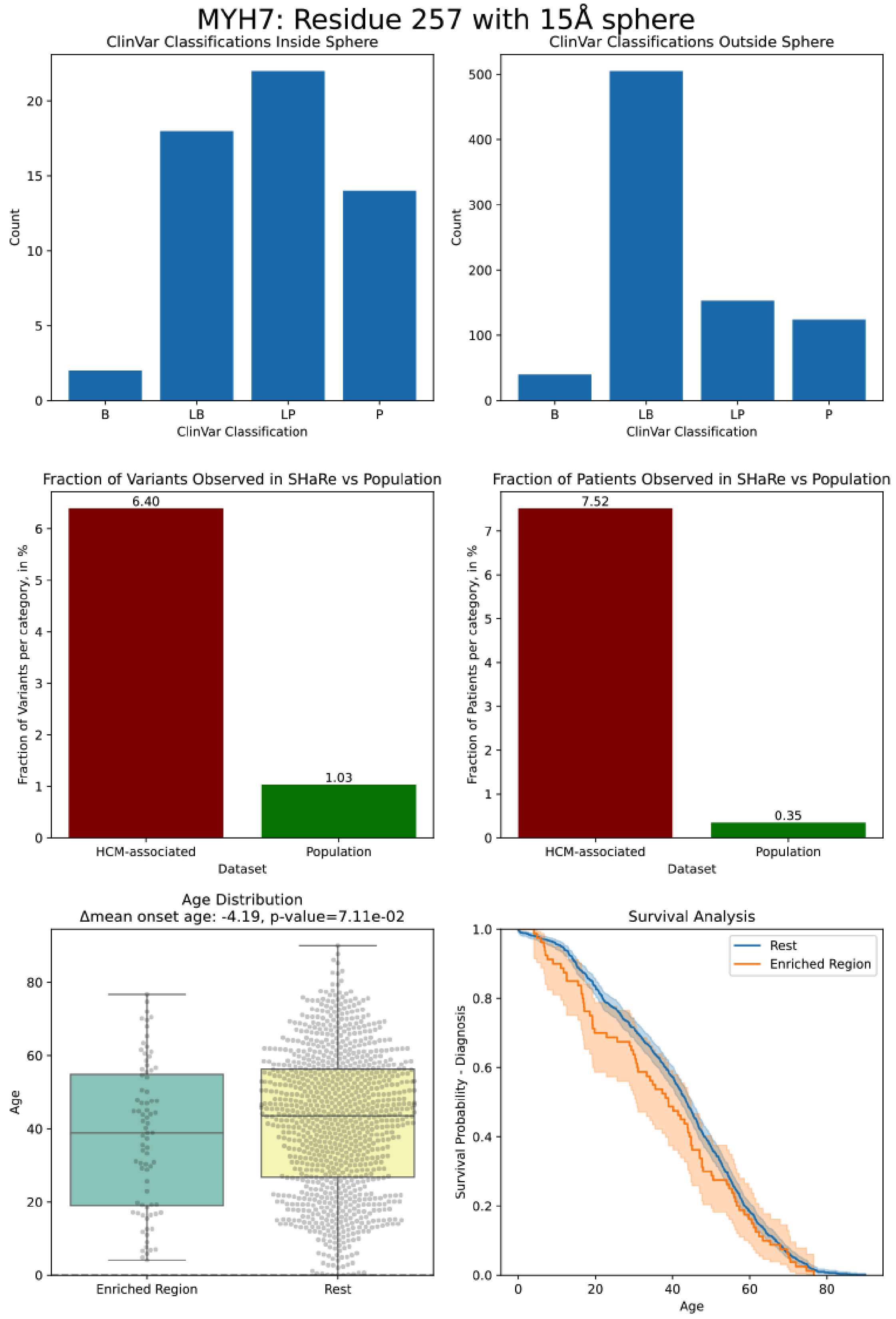


***Figure S7.*** *Validation plots for MYH7 transducer domain cluster located on residue 257*

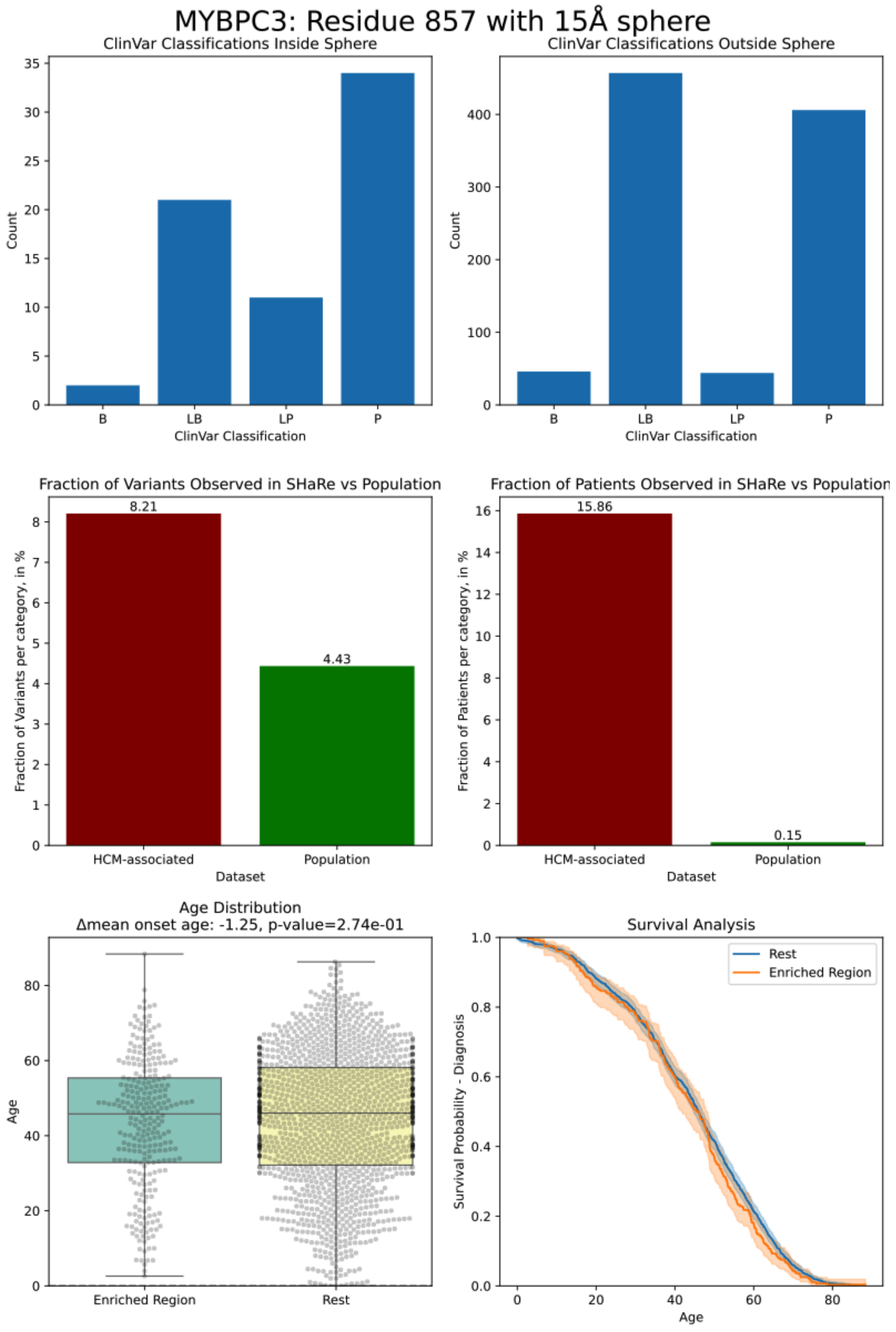


***Figure S8.*** *Validation plots for MYBPC3 C6  domain cluster located on residue 857*

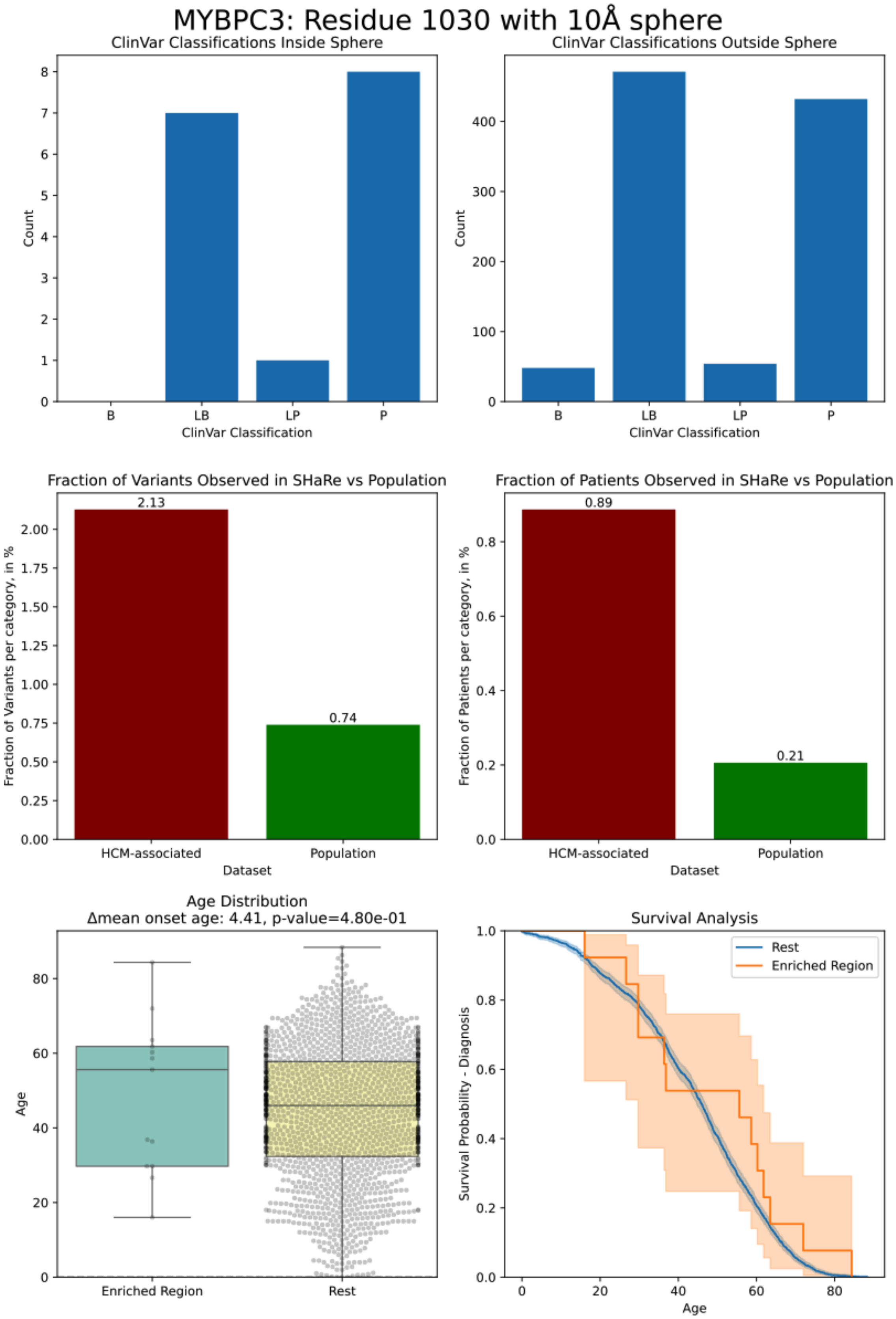
MYBPC3: Residue 1030 with 10Å sphere
ClinVar Classifications Inside Sphere
Count
ClinVar Classification
B
LB
LP
P
ClinVar Classifications Outside Sphere
Count
ClinVar Classification
B
LB
LP
P
Fraction of Variants Observed in SHaRe vs Population
Fraction of Variants per category, in %
2.13
0.74
HCM-associated
Population
Dataset
Fraction of Patients Observed in SHaRe vs Population
Fraction of Patients per category, in %
0.89
0.21
HCM-associated
Population
Dataset
Age Distribution
Δmean onset age: 4.41, p-value=4.80e-01
Age
Enriched Region
Rest
Survival Analysis
Survival Probability - Diagnosis
Rest
Enriched Region
Age

***Figure S9.*** *Validation plots for MYBPC3 C8  domain cluster located on residue 1030*

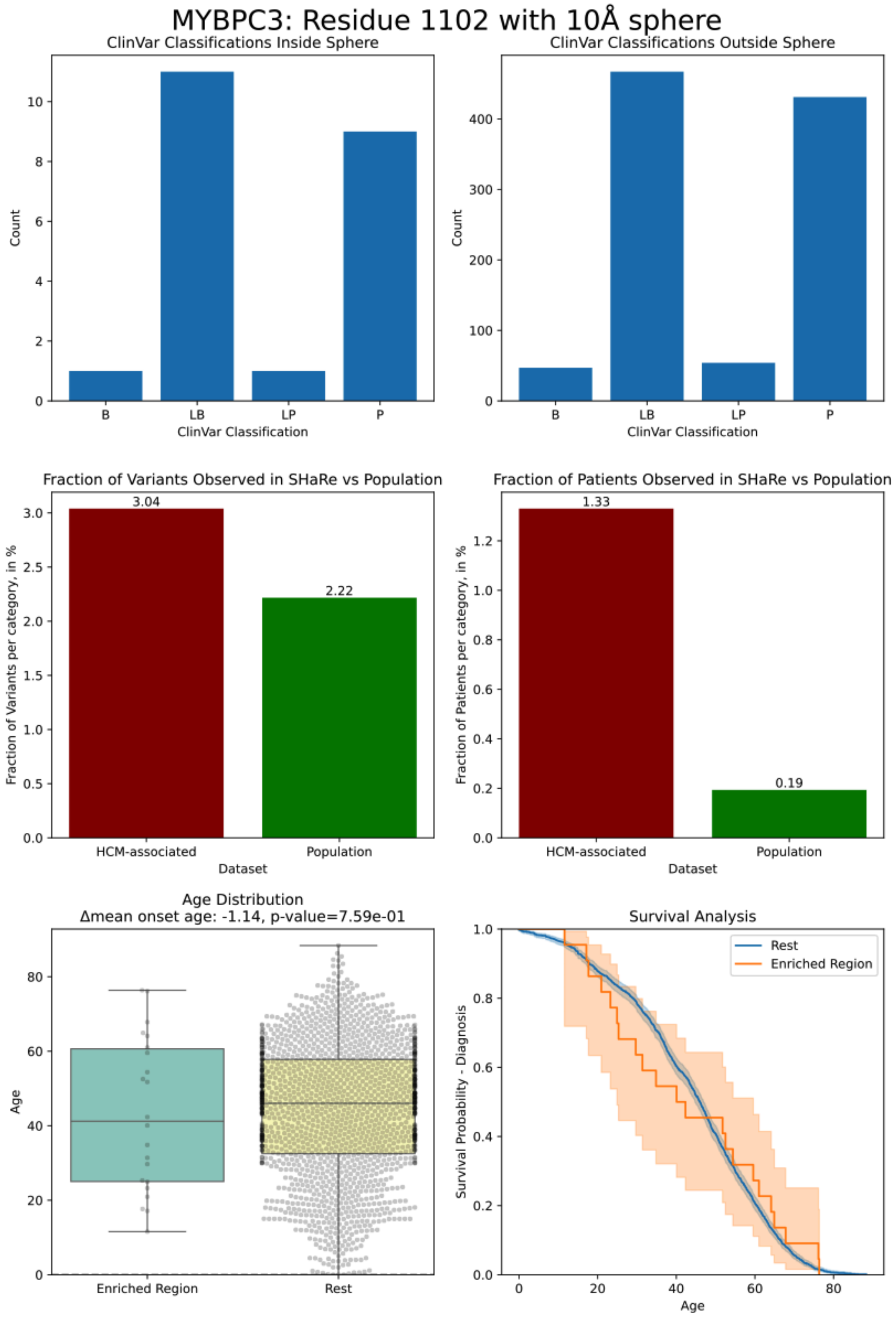

***Figure S10.*** *Validation plots for MYBPC3 C9  domain cluster located on residue 1102*

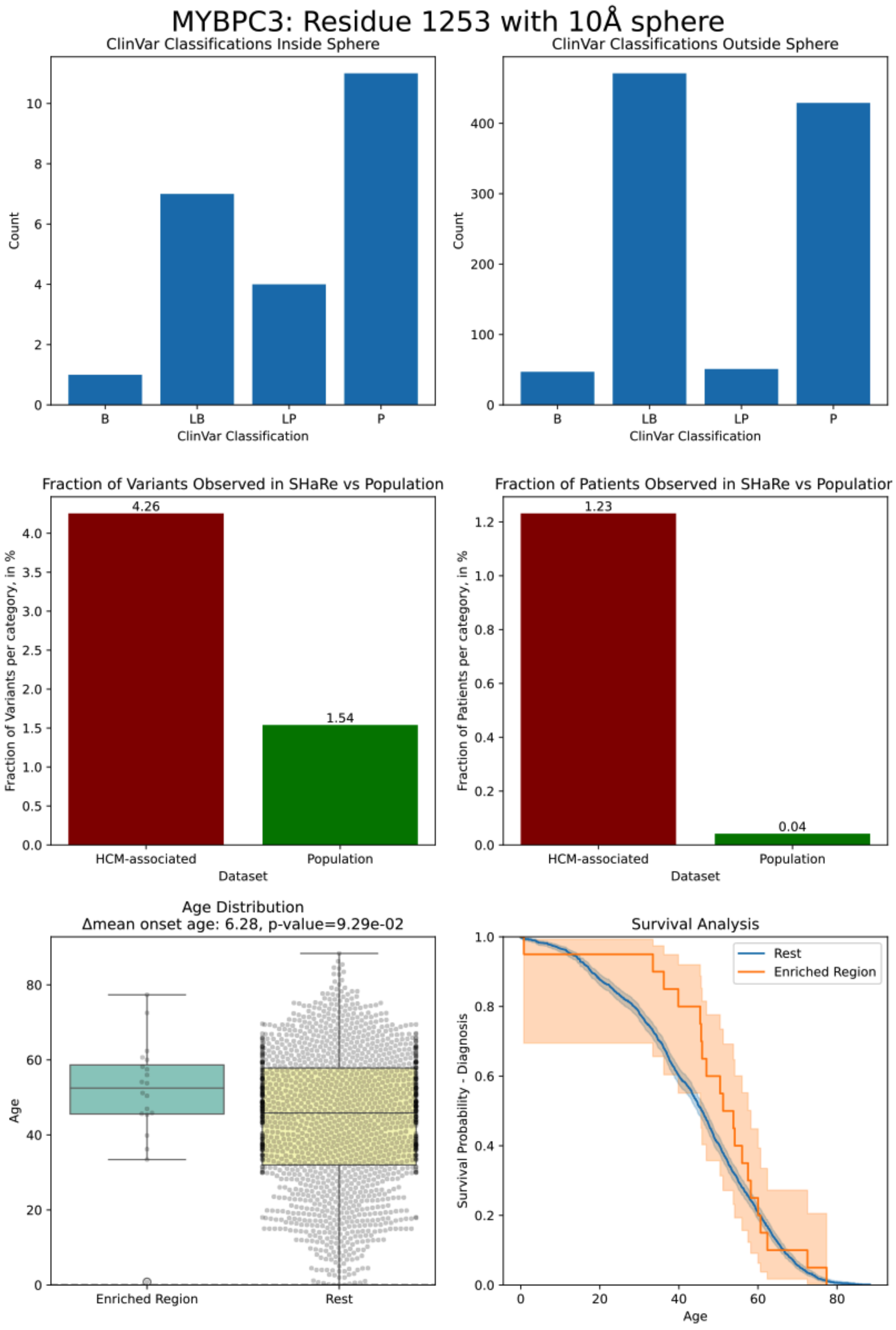

***Figure S11.*** *Validation plots for MYBPC3 C10  domain cluster located on residue 1253*

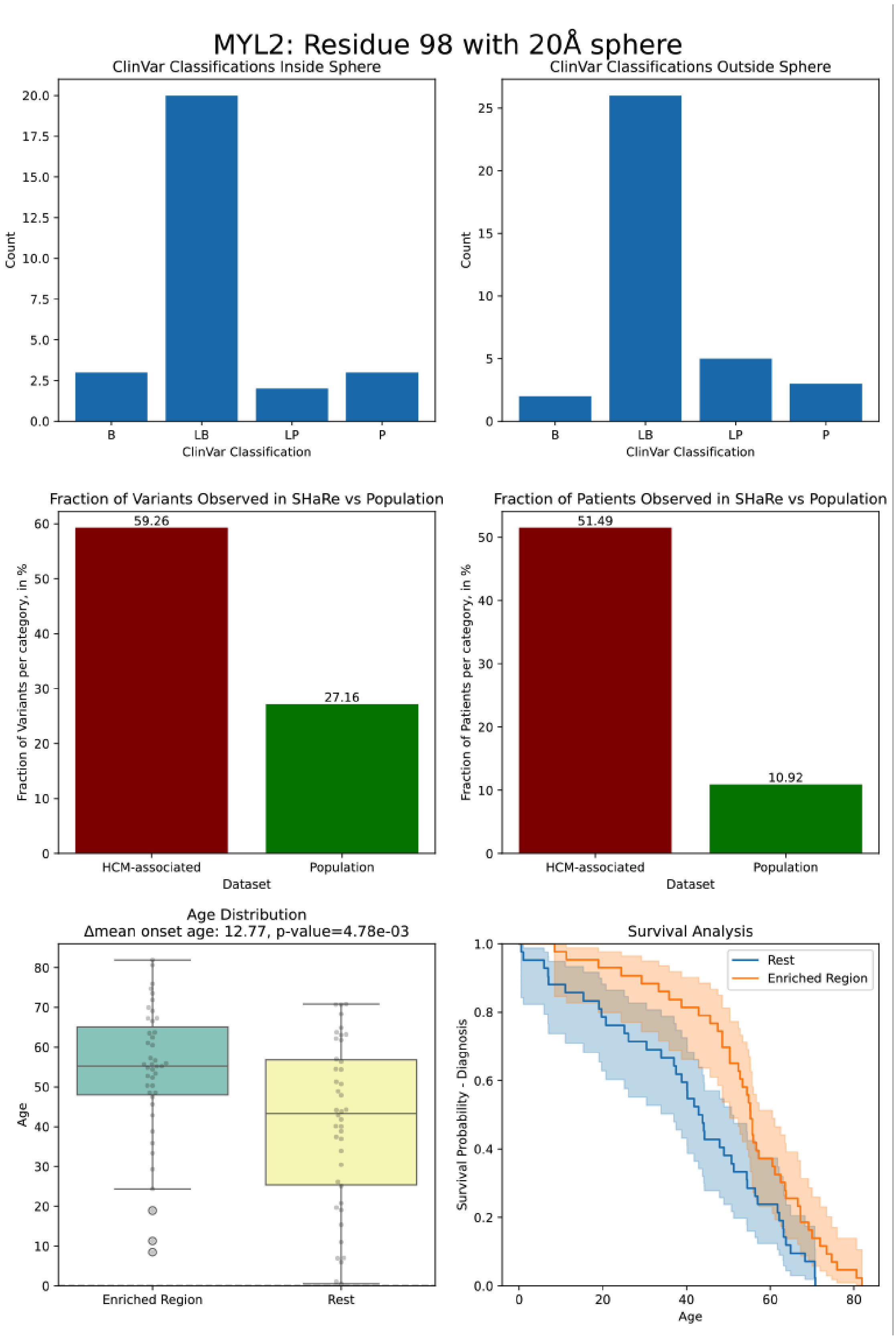

***Figure S12.*** *Validation plots for MYL2 cluster surrounding the MYH7 lever arm located on residue 98*

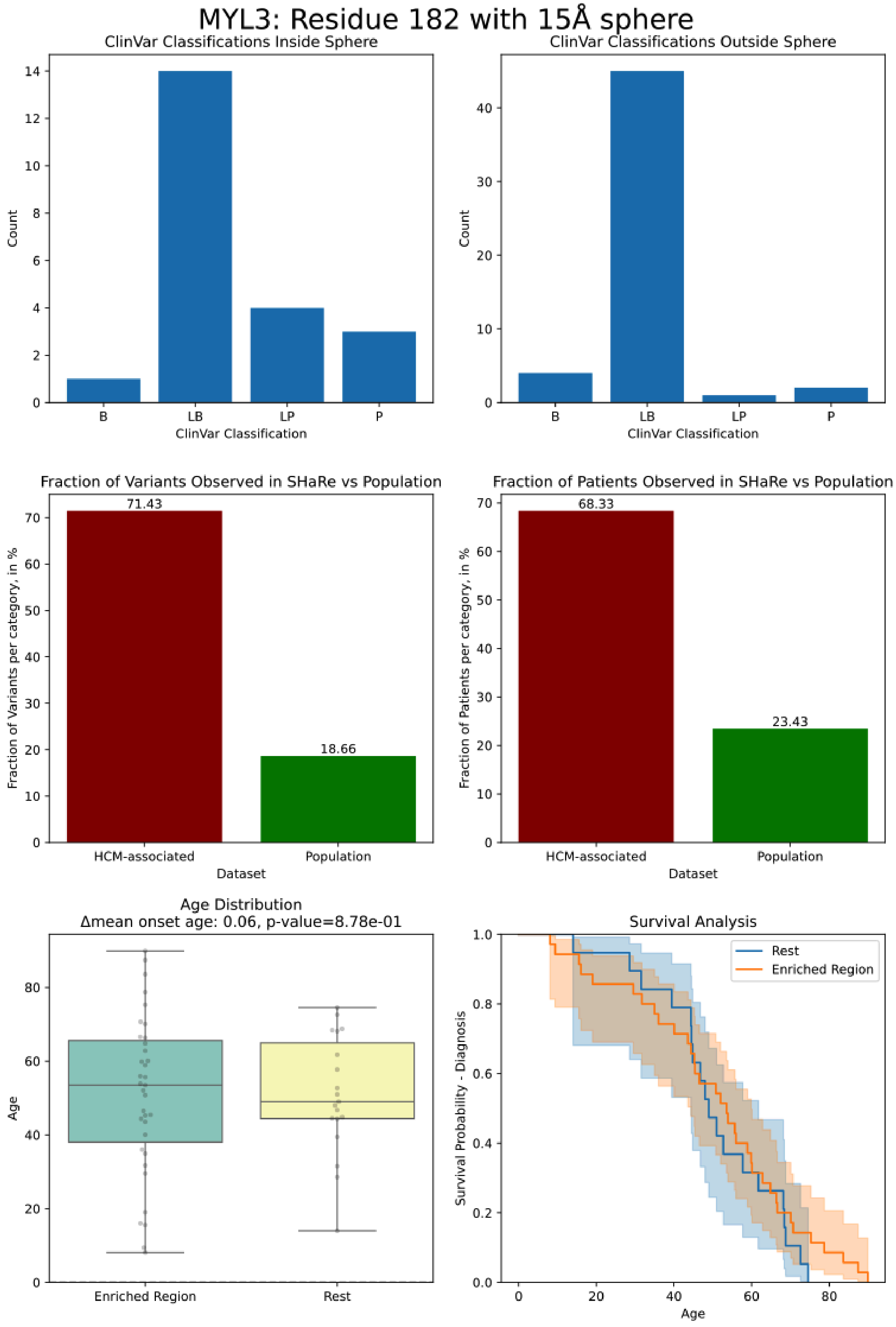


***Figure S13.*** *Validation plots for MYL3 EF-hand 2 cluster located on residue 182*

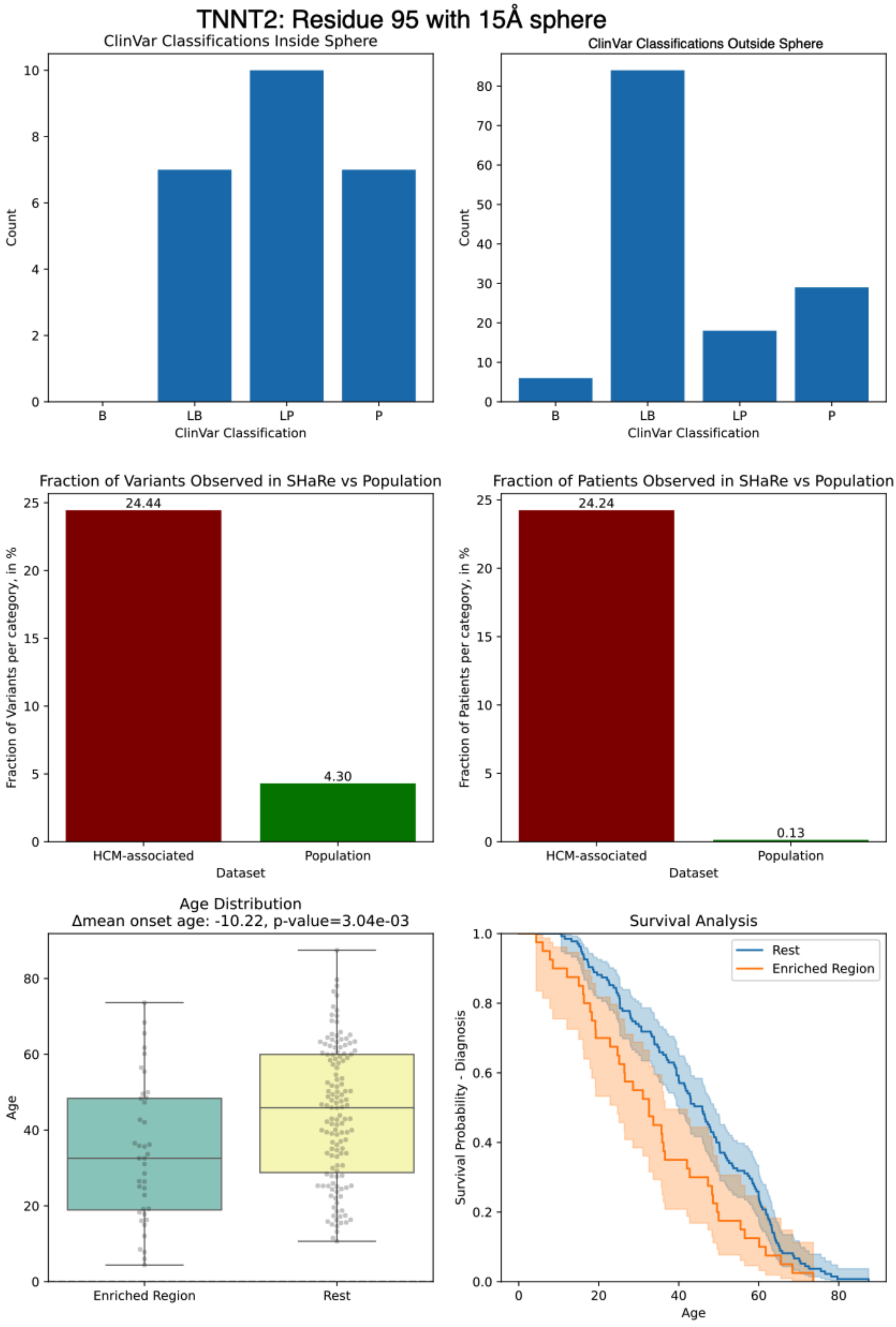


***Figure S14.*** *Validation plots for TNNT2 tropomyosin-actin binding site cluster located on residue 95*

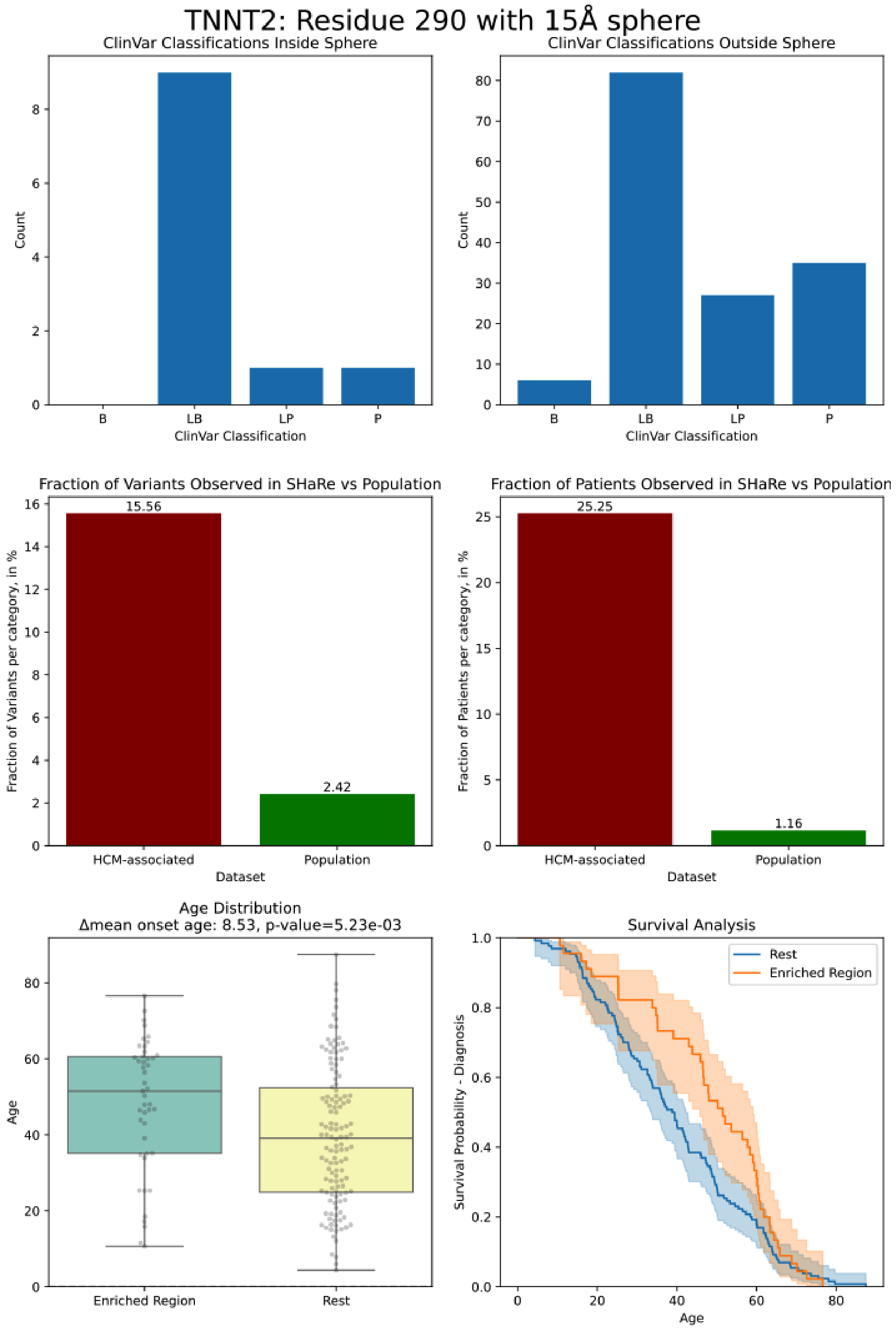


***Figure S15.*** *Validation plots for TNNT2 C-terminus cluster located on residue 290*

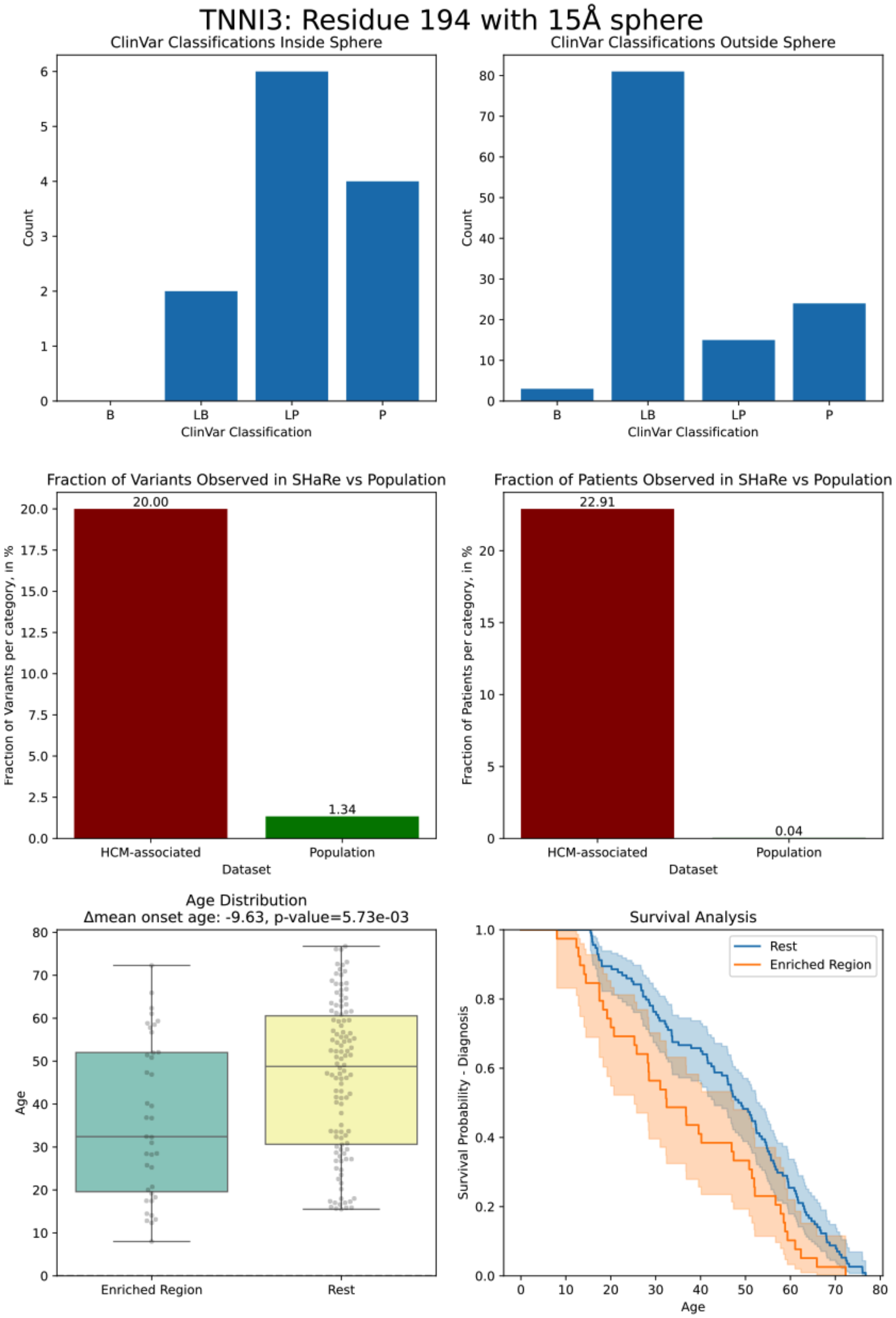


***Figure S16.*** *Validation plots for TNNI3 mobile domain cluster located on residue 194*

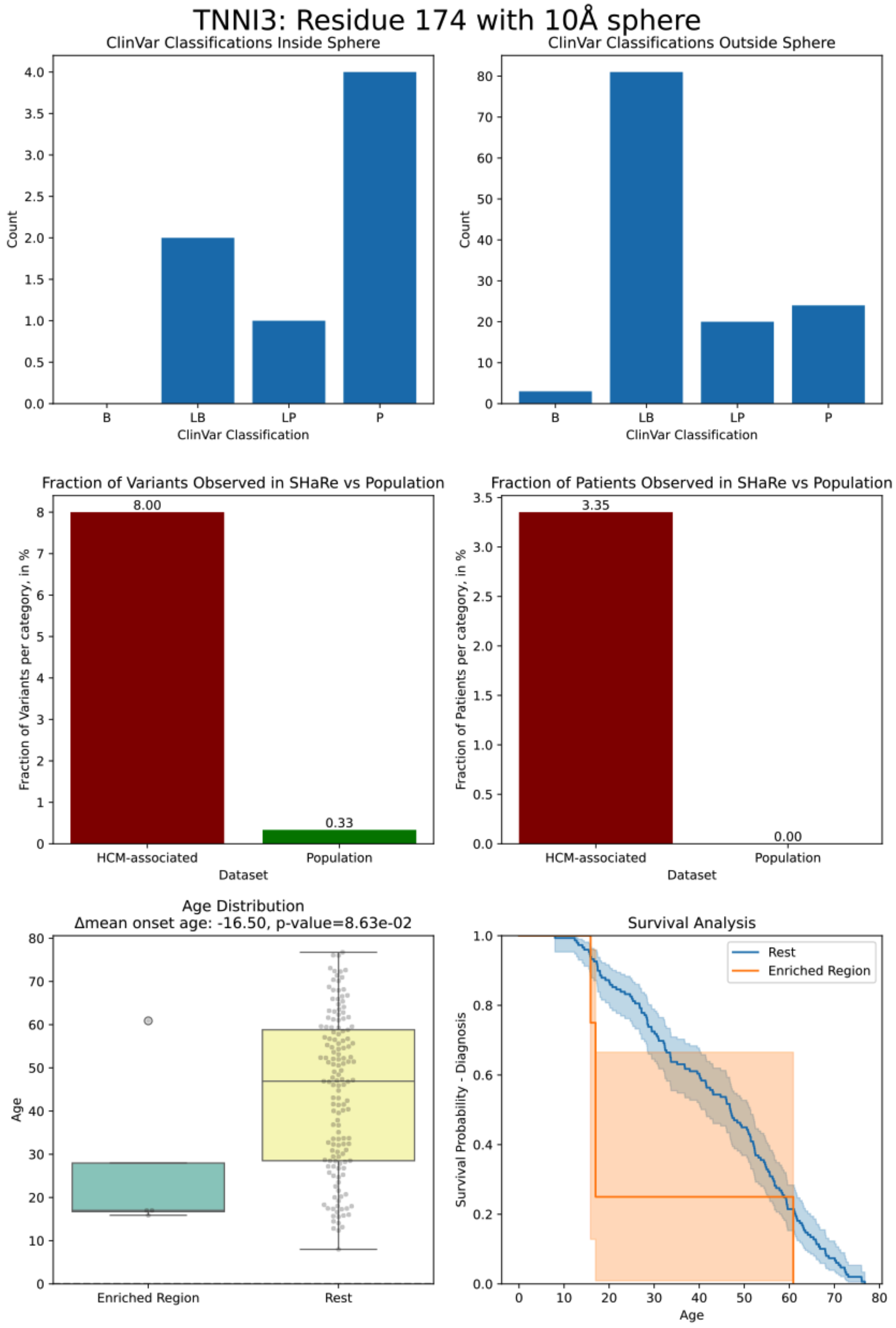


***Figure S17.*** *Validation plots for TNNI3 cluster connecting mobile domain to switch domain located on residue 174*

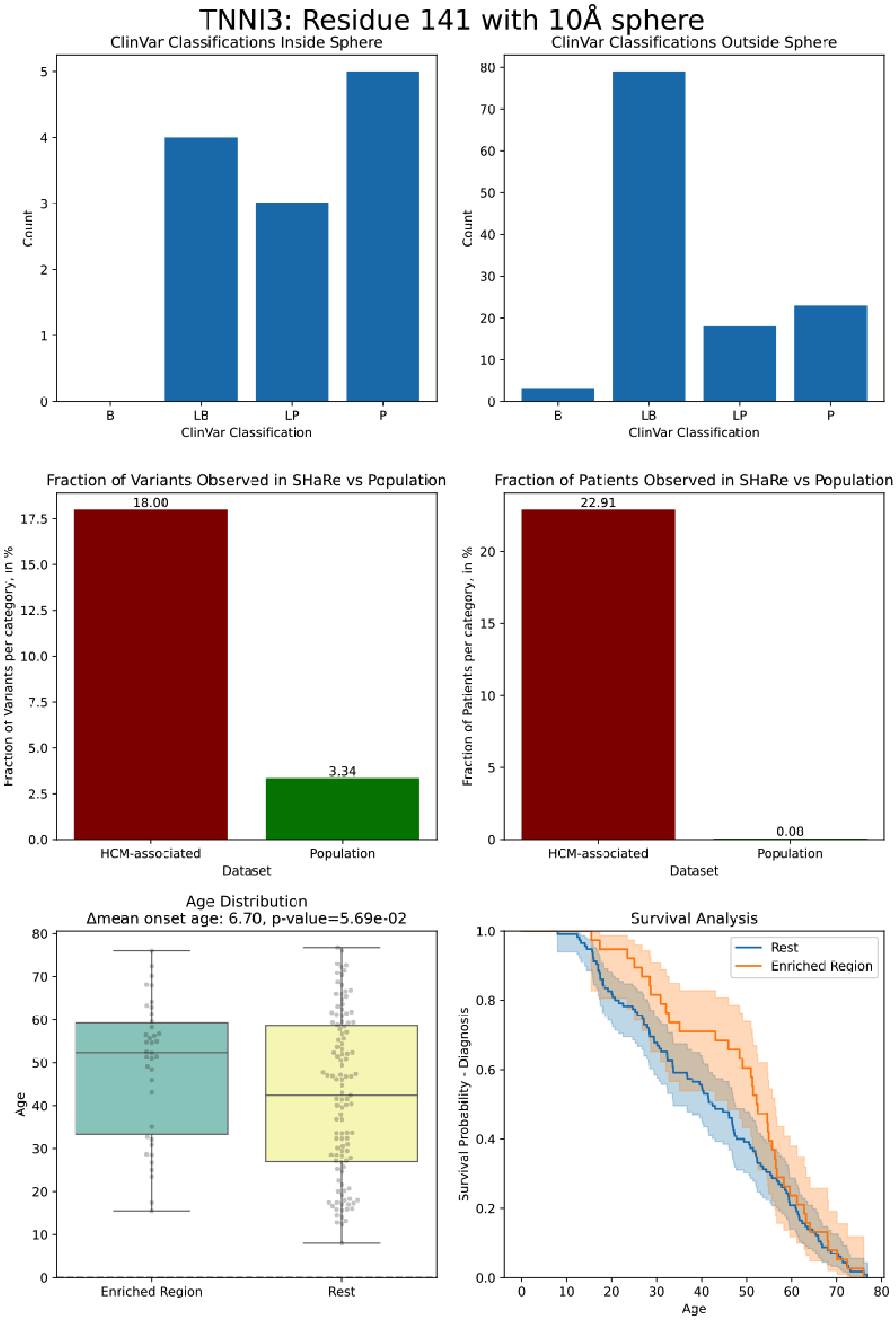


***Figure S18.*** *Validation plots for TNNI3 switch domain cluster located on residue 141*

#### Tables

| Gene | Domain of center residue | Expected # variants in sphere | Observed # variants in sphere |
|---|---|---|---|
| *MYH7* | Converter | 2.2 | 8 |
| *MYH7* | Relay Helix / Myosin mesa (residue 493) | 1.2 | 1 |
| *MYH7* | SH1-SH2 Helix / Myosin mesa (residue 696) | 1.1 | 2 |
| *MYH7* | Transducer / Near nucleotide-binding site | 2.0 | 3 |
| *MYBPC3* | C6 - interacts with the myosin tail | 4.9 | 5 |
| *MYBPC3* | C8 - interacts with the IHM of myosin | 1.7 | 2 |
| *MYBPC3* | C9 - interacts with the myosin tail | 2.2 | 2 |
| *MYBPC3* | C10 - interacts with the IHM of myosin | 2.4 | 3 |
| *MYL2* | Surrounding *MYH7* lever arm | 1.9 | 3 |
| *MYL3* | EF-hand 2, surrounding *MYH7* lever arm | 2.8 | 8 |
| *TNNT2* | Tropomyosin-actin-binding site | 1.0 | 3 |
| *TNNT2* | C-terminus | 0.8 | 4 |

| | | | |
|---|---|---|---|
| *TNNI3* | Mobile domain | 0.6 | 0 |
| *TNNI3* | Connecting Switch to mobile domain | 1.0 | 1 |
| *TNNI3* | Switch domain | 0.9 | 3 |

***Table S1.*** *Comparison of expected vs. observed number of disease-associated variants within identified regions*
